# Third-party prosocial behavior in adult female rats is impaired after perinatal fluoxetine exposure

**DOI:** 10.1101/763276

**Authors:** Indrek Heinla, Roy Heijkoop, Danielle J. Houwing, Jocelien D.A. Olivier, Eelke M.S. Snoeren

**Author notes:** **Correspondence** Eelke Snoeren, Department of Psychology, UiT the Arctic University of Norway, Postboks 6050 Langnes, 9037 Tromsø, Norway.

## Abstract

SSRIs are commonly used to treat pregnant women with depression. However, SSRIs can cross the placenta and affect the development of the fetus. The effects of perinatal SSRI exposure, and especially the effects on social behavior, are still incompletely documented. This study first aims to investigate whether rats show prosocial behavior in the form of consolation behavior. Secondly, it aims to investigate whether perinatal SSRI exposure affects this prosocial behavior. At last, we investigate whether the behavior changed after the rats had been exposed to an additional white-noise stressor.

Rat dams received 10 mg/kg/d fluoxetine (FLX) or vehicle (CTR) via oral gavage from gestational day 1 until postnatal day 21. At adulthood, the rat offspring were housed in four cohorts of 4 females and 4 males in a seminatural environment. As prosocial behaviors are more prominent after stressful situations, we investigated the behavioral response of rats immediately after natural aggressive encounters (fights). Additionally, we studied whether a stressful white-noise exposure would alter this response to the aggressive encounters.

Our study indicates that CTR-female rats are able to show third party prosocial behavior in response to witnessing aggressive encounters between conspecifics in a seminatural environment. In addition, we showed that perinatal FLX exposure impairs the display of prosocial behavior in female rats. Moreover, we found no signs of prosocial behavior in CTR- and FLX-males after natural aggressive encounters. After white-noise exposure the effects in third party prosocial behavior of CTR-females ceased to exist. We conclude that female rats are able to show prosocial behavior, possibly in the form of consolation behavior. In addition, the negative effects of perinatal fluoxetine exposure on prosocial behavior could provide additional evidence that SSRI treatment during pregnancy could contribute to the risk for social impairments in the offspring.

## 1. Introduction

Selective serotonin reuptake inhibitors (SSRIs) are the most prevalent treatment for women with depression during pregnancy [1, 2]. Nonetheless, since SSRIs can cross the placenta and appear in breastmilk [3–5], the question rises how this treatment might affect the developing fetus. SSRI exposure during development affects the serotonergic system in the fetus [5, 6]. While serotonin acts as neurotransmitter at adulthood, it is a neurotrophic factor during early brain development. More specifically, perinatal SSRI exposure can affect the regulation of cell division, differentiation, migration, and synaptogenesis [7, 8]. Whether or not these effects also have consequences later in life remains unclear, but it is assumed that perinatal SSRI exposure affects the serotonergic function and vulnerability to affective disorders [9].

Several studies have found an association between perinatal SSRI exposure and disturbed sleep patterns, affected social-emotional development, and increased internalizing and externalizing behavior in the offspring [10–12]. Recently, an ongoing debate started about whether children whose mothers were treated with SSRIs during pregnancy have an increased risk to develop autism spectrum disorder (ASD). Some studies show a clear correlation between SSRI treatment and increased odds for ASD in the offspring [13–17], whereas others do not find this link or suggest that this increased risk is caused by the depression itself rather than the SSRIs [18, 19]. The mothers who do take antidepressants during pregnancy most likely suffer from a more severe depression, and untreated depression during pregnancy may also have negative impact on the offspring [18, 20–23]. In fact, when controlled for maternal mood and stress, the link between antenatal SSRI use and the occurrence of ASD in the offspring does not persist [24].

Where human epidemiological studies struggle with the lack of control over variables and possible confounding factors, animal studies can provide a more fundamental insight into underlying mechanisms of illnesses and treatments. By treating healthy mothers with antidepressants, one can discern the effects of the drug exposure during pregnancy on neurodevelopmental alterations in the offspring.

Although the link with ASD and perinatal SSRI exposure remains controversial, effects of perinatal SSRI exposure on social behavior have been found. For example, some studies showed that both pre- and early postnatal SSRI treatment can decrease social play behavior and communicative skills (ultrasonic vocalizations) in young rodent offspring [25–29], with the exception of one study that showed an increase in social play [30]. In terms of social interaction at adulthood, the findings are more controversial. Some studies have indicated that developmental SSRI exposure decreases social interaction in either male offspring [25, 27] or female offspring [31], while others found an increase in sniffing behavior towards conspecifics [32, 33]. When looking at the motivation for social interaction in particular, the majority found that SSRI exposure negatively affects the motivation to start social contact [26, 28, 29, 34]. The inconsistent results make it impossible to draw conclusions about the risk for long-term affected social behavior in the offspring, just as it did not take the role of passive social behavior into account. Recently, we found that both male and female rats that had been perinatally exposed to fluoxetine more often spent passive moments in the company of a conspecific (social resting) compared with control rats [31], confirming the risk for affected social behavior resulting from perinatal SSRI exposure.

Interestingly, another important element of social behavior which received less attention in research is *prosocial behavior*. Prosocial behavior is defined as behavior that is intended to benefit another individual in distress, and includes helping and consolation behavior [35]. The difference between helping and consolation behavior is that consolation is an increase in affiliative contact (like grooming and hugging) in response to and directed toward a distressed individual by a bystander, which produces a calming effect [36]. Helping behavior, on the other hand, also improves the status quo of another individual, but does not require direct contact.

With regard to helping behavior, several studies have been performed in rats. Rats do liberate conspecifics trapped in restrainers or soaked in water arenas, even when there is no social reward at the end of the test [37–39]. When rats were placed in a food-foraging task, they behave prosocially by choosing the option that provides their cage mate with food as well [40]. Until now, consolation behavior has been documented in several species (including great apes, dogs, wolves, rooks, elephants, and prairie voles), mainly in the context of naturally occurring aggressive conflicts [36, 41–46]. However, no studies have yet confirmed that rats show consolation behavior, especially not in a more natural setting.

This study, therefore, first aims to investigate whether rats show prosocial behavior in the form of consolation behavior. The second aim of the study was to investigate whether perinatal SSRI exposure affects this prosocial behavior. At last, we investigate whether the behavior changed after the rats had been exposed to an additional white-noise stressor. We performed an observational study in which we used a seminatural environment in which cohorts of eight rats are group housed for 8 days. The advantage of this environment is that the rats are able to express their full repertoire of natural behaviors. If consolation behavior exists in rats, it is expected to happen after stressful events like naturally occurring aggressive encounters between conspecifics. We therefore observed the behavior of all rats during the 15 minutes after fights and expected to find an increase in active social behavior directed at conspecifics. In a previous study, we found that phenotypes in perinatally SSRI exposed offspring can change after experiencing a stressful event [31]. We therefore also investigated whether the performance of prosocial behavior is changed after an additional stressor.

## 2. Materials and methods

This study describes the data from subsequent analyses of video recordings from another study. This means that the rats and thus the procedures were the same as mentioned in [31], but the behavioral observations were chosen and designed specifically for the purpose of the current study. The materials and methods describe the steps that were required for the current study.

### 2.1 Housing conditions

Male (n=10) and female Wistar rats (n=10), weighing 200-250g on arrival, were obtained from Charles River (Sulzfeld, Germany). They were used as dams and potential fathers of the offspring, and housed in same sex pairs in Makrolon IV cages (60 × 38 × 20 cm) under a 12 : 12 h reversed light/dark cycle (lights on 23.00 h) at 21 ± 1 °C and 55 ± 10% relative humidity. Standard rodent food pellets (standard chow, Special Diets Services, Witham, Essex, UK) and tap water were available ad libitum, and nesting material was present.

For the mating process, each female was housed with one male in a Makrolon IV cage for 24 hours, after which she returned to her same sex pair for the following two weeks of gestation. On gestational day 14, the females were single-housed in Makrolon IV cages for delivery, in which they stayed until weaning of the pups.

The offspring were housed together with their mother until weaning (postnatal day 21). Thereafter until the start of the experiment, the offspring were housed in groups of two/three same sex littermates in Makrolon IV cages. Ears were punched for individual recognition. The animals were left undisturbed except during cage cleaning.

All experimentation was carried out in agreement with the European Union council directive 2010/63/EU. The protocol was approved by the National Animal Research Authority in Norway.

### 2.2 Breeding and fluoxetine treatment

Before mating, the dams were checked daily for the expression of lordosis behavior upon male rat mounting attempts by placing the females together with a male rat for a maximum of 5 minutes. As soon as lordosis was observed, the dam was considered to be in proestrus, and mating could be started. For the mating process, each female in proestrus was housed with one male for approximately 24 hours (Gestational day 0) to induce pregnancy.

From gestational day 1 (G1, the day after mating) until postnatal day 21 (PND21), a total of 6 weeks, the mothers of the experimental animals returned to their home cages and were treated daily with a stainless steel feeding needle per oral gavage with either fluoxetine (10 mg/kg, n= 6 dams) or vehicle (Methylcellulose 1%, (Sigma, St. Louis, MO, USA), n=4 dams) in a volume of 5 ml/kg. Fluoxetine pills were crushed and dissolved in sterile water (2 mg/ml), while 1% methylcellulose was dissolved directly in sterile water. The females were weighed every three days to determine the dose for the following three days. The dose of fluoxetine was based on comparison to human treatment [27, 47]. Near the end of pregnancy, dams were checked twice per day (9 p.m. and 15 p.m.) for delivery. Three females did not become pregnant (2 dams treated with fluoxetine, and 2 dam treated with vehicle) and one female (fluoxetine treatment) delivered dead offspring, therefore no offspring of these dam were included in the study, reducing the number to n=3 dams per treatment. An overview of the whole procedure from the beginning of antidepressant treatment until the end of testing of the offspring is presented in Figure 1.

**Figure 1.**
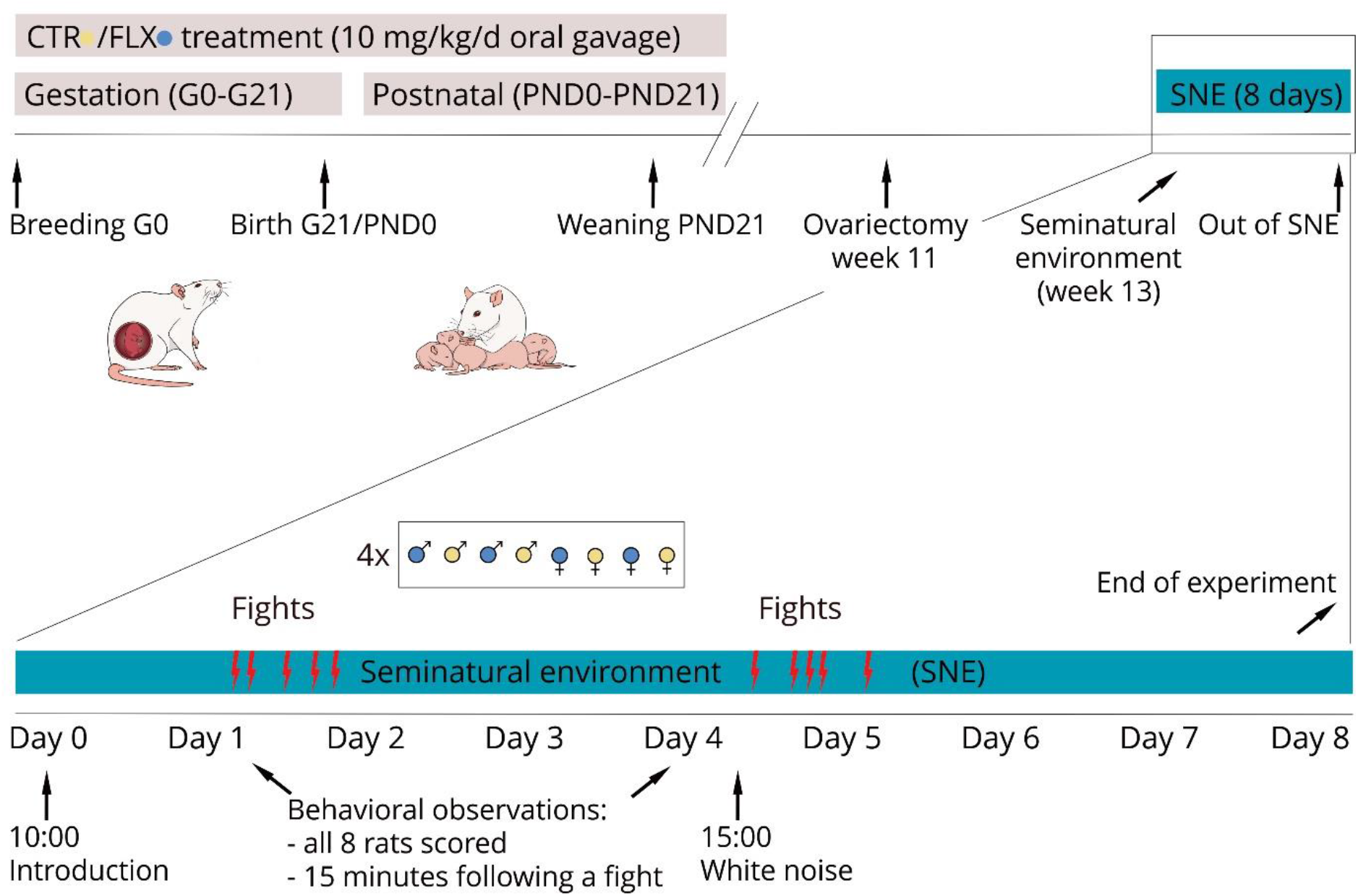
Timeline of the experiment. Pregnant rats were treated via oral gavage with either vehicle (CTR) or 10 mg/kg/d fluoxetine (FLX) from gestational day 0 (G0) until weaning of the pups (PND21). The behavior of the offspring at adulthood was evaluated in the seminatural environment (SNE) after naturally occurring aggressive encounters (fights). An additional stressor, a 10-minute white-noise exposure, was presented in day 4, enabling the comparison between pre- and post-stress behavior.

### 2.3 Design of the study

After birth, litters were not culled. Pups were weaned at PND 21 and housed in groups of two or three same sex littermates in Makrolon IV cages (see Table S2 for more details), and left undisturbed until an age of 13-18 weeks (adulthood). From these offspring, 32 rats (of the 6 litters) were distributed in 4 cohorts and used for behavioral evaluation in the seminatural environment. A cohort of rats consisted of two male offspring from control mothers (CTR-males), two males from fluoxetine-treated mothers (FLX-males), two females from control mothers (CTR-females) and two females from fluoxetine-treated mothers (FLX-females). The total of 4 cohorts thus resulted in n=8 per treatment and sex group for data analysis.

Within a cohort, same sex rats came from different litters and were therefore unfamiliar to each other. However, due to a limited amount of litters available, some animals had one sibling from the opposite sex in the same cohort. These littermates had been housed in different home cages since weaning. Details of the litter distribution in the cohorts can be found in the supplemental materials of [31].

For the purpose of the previous study [31], the females had undergone ovariectomy two weeks before entering the seminatural environment (see 2.5 for description of this environment). For the current study this means that all females were ovariectomized and without hormonal priming, which has the benefit that sexual behavior did not occur and thus did not influence our data.

### 2.4 Procedure in the seminatural environment

The day before the subjects were introduced into the seminatural environment, they were shaved on the back and tail-marked under isoflurane anesthesia for individual recognition. (For more details, see [31]) In addition, the offspring were weighed and no differences in body weight were found between CTR and FLX animals.

Each cohort of rats was introduced into the seminatural environment on the first day (Day 0) at 10 a.m., and removed at the end of the experiment on day 8 at 10 a.m. Between cohorts, the seminatural environment was thoroughly cleaned to remove olfactory cues from previous animals.

### 2.5 Description of the seminatural environment

The seminatural environment (2.4 × 2.1 × 0.75 meters) setup is previously described in [31, 48, 49]. It consists of a burrow system and an open field area, which are connected by four 8 × 8 cm openings. The burrow system consists of an interconnected tunnel maze (7.6 cm wide and 8 cm high) with 4 nest boxes (20 × 20 × 20 cm), and is covered with Plexiglas. The open area is an open area with 0.75 meter high walls, and contains two partitions (40 × 75 cm) to simulate obstacles in nature. A curtain between the burrow and the open field allowed the light intensity for both arenas to be controlled separately. The burrow system remained in total darkness for the duration of the experiment, while a day-night cycle was simulated in the open area with a lamp 2.5 m above the center that provided 180 lux from 22.45h to 10.30h (the equivalent of daylight) and approximately 1 lux from 10.30h to 11.00h (the equivalent of full moonlight). The light gradually increased/decreased during 30 minutes between 1 and 180 lux.

The floors of both the open area and on the burrow system were covered with a 2 cm layer of aspen wood chip bedding (Tapvei, Harjumaa, Estonia). In addition, the nest boxes were provided with 6 squares of nesting material each (nonwoven hemp fibres, 5 × 5 cm, 0.5 cm thick, Datesend, Manchester, UK), and in the open area 3 red polycarbonate shelters (15 × 16.5 × 8.5 cm, Datesend, Manchester, UK) were placed and 12 aspen wooden sticks (2 × 2 × 10 cm, Tapvei, Harjumaa, Estonia) were randomly distributed. Food was provided in one large pile of approximately 2 kg in the open area close to the water supply. Water was available ad libitum in 4 water bottles.

Two video cameras (Basler) were mounted on the ceiling 2 meter above the seminatural environment: one above the open field and an infrared video camera above the burrow system. Videos were recorded using Media recorder 2.5 by direct connection to a computer allowing the data to be stored immediately on an external hard drive. Every 24 hours, the recording was stopped manually and restarted to create recordings with a length of 24h. This was done to assure that if a recording error would occur, data from only one day would be lost.

### 2.6 White-noise

To investigate the response of the offspring to a stressful event, and compare the behavior before and after, the rats were exposed to 90dB white-noise containing all frequencies at the same time, produced by a white-noise generator (Lafayette instruments, Lafayette, IN). This white-noise was played to the rats via two loudspeakers (Scan-Speak Discovery 10F/8414G10, HiFi Kit Electronic, Stockholm), one of which was placed in the open field and the other in the burrow area. Exposure occurred on day 4 at 15.00h and lasted for 10 minutes.

### 2.7 Behavioral observations

In order to find potential prosocial behavior after naturally occurring stressful situations, the video recordings were screened for aggressive encounters between the animals. An aggressive encounter was registered when two rats attempted to bite and/or wrestle each other aggressively, usually accompanied by loud high-pitch screeching, resulting in one animal (loser) fleeing the area. Boxing, nose-off and playful wrestling were not considered aggressive encounters.

For each aggressive encounter, the role of each rat during this encounter (*role in fight*) was determined with 4 possible options: 1) winner - chasing the conspecific after the encounter; 2) loser - running away, escaping from the winner; 3) witness - was in immediate vicinity of the fight and/or paid attention to the ongoing fight (by facing the fight) or 4) non-witness - was not in immediate vicinity of the fight and did not pay attention to the ongoing fight. The aggressive encounters happened spontaneously and were randomly selected upon sequence. The majority of fights were between two male rats, except for 2 female versus female fights, and 2 fights where the winner was a male and the loser a female.

The behavior of each cohort member was observed during 15 minutes immediately after each aggressive encounter. Originally, we planned to include aggressive encounters at two different time points: A) five fights at baseline (the pre-stress condition, starting on day 1 of the experiment) and B) five fights after the white-noise exposure (the post-stress condition, starting immediately after the white-noise exposure on day 4). However, in order to test our hypothesis, the most important comparison is to investigate the behavior of each rat in instances when they witnessed aggressive encounters versus instances when they did not. In order to assure that most of the rats were tested as witness *and* non-witness, we continued the search for aggressive encounters until most rats had played both roles at least once. This resulted in an outcome of 5-6 fights per rat at baseline, and 5-7 fights in post-stress. However, some rats never played both roles during the encounters, which resulted in a few missing data points.

The duration and/or the frequency of the behaviors defined in table 1a was registered by an observer, blind for the treatment of the animals (and therefore also blind for the expected outcome on behaviors), in the Observer XT software (Noldus Information Technology, The Netherlands; version 12.5). In addition, we registered towards which conspecific social and conflict behaviors were directed, and in what location (open area, tunnels or nest box) the behaviors took place.

**Table 1a.** Description of recorded behaviors

| *Behavior* | *Description* |
| --- | --- |
| *Allogrooming* | Subject is grooming another rat |
| *Sniffing others* | Subject is sniffing another rat's face or body area |
| *Sniffing anogenitally* | Subject is sniffing another rat's anogenital area |
| *Self-grooming alone* | Subject grooms itself |
| *Self-grooming near others* | Subject grooms itself in close proximity of others |
| *Walking* | Subject is moving around in the environment |
| *Walking over/under other* | Subject is walking over/under other rat |
| *Resting/immobile* | Subject is sleeping or standing still with no or minimal head movement |
| *Resting with others* | Subject is huddling/resting in the close vicinity of others |
| *Non-social exploration* | Subject is actively sniffing around and/or rearing in the environment |
| *Digging/carrying* | Subject digs bedding material or moves around nesting material or food pellets |
| *In opening facing open area* | Subject is staying in the opening between open area and the burrow and looking towards open area |
| *In opening facing burrow* | Subject is staying in the opening between open area and the burrow, looking towards the burrow |
| *Fighting* | Subject is fighting with another rat |
| *Nose-off* | Subject is facing another rat and aggressively posturing towards it |
| *Defensive posture* | Subject is defensively posturing near other rats |
| *Boxing* | Subject is facing another rat and frequently hitting it with front paws |
| *Wrestling* | Subject wrestles with another rat |
| *Fleeing* | Subject is fleeing from another rat |
| *Chasing* | Subject chases other rat after a conflict situation |
| *Drinking* | Subject drinks |
| *Witnessing* | Was in immediate vicinity of the fight and/or paid attention to the ongoing fight |
| *Non witnessing* | Was not in immediate vicinity of the fight and did not pay attention to the ongoing fight |

### 2.8 Statistical analysis

As shown in Table 1b, behavioral clusters were created for behaviors that fit in the same behavioral group. For each rat, we calculated the frequency and the total time spent on each behavior (and behavioral cluster) after each aggressive encounter. In addition, we calculated the frequencies and durations of behaviors during all encounters taken from the perspective of the role the rat played during the fights (witness versus non-witness). This analysis was performed for both pre- and post-stress aggressive encounters.

**Table 1b.** Description of behavioral clusters

| *Cluster* | *Behaviors within the cluster* |
| --- | --- |
| *Active social behavior* | Allogrooming, Sniffing others, Sniffing anogenitally |
| *Passive social behavior* | Resting with others |
| *Social context* | Allogrooming, Sniffing others, Sniffing anogenitally, Self-grooming near others, Resting with others |
| *General activity* | Walking, Walking over/under other, Non-social exploration |
| *General passive* | Resting/immobile, Resting with others |
| *Conflict* | Fighting, Nose-off, Defensive posture, Kicking, Boxing, Wrestling, Fleeing, Chasing |
| *Self-grooming* | Self-grooming alone, Self-grooming near others |

A Shapiro–Wilk test showed no homogeneity of variance. However, to control for multiple comparisons, a linear mixed model design was used first which included all fights as repeated measure, and the conditions *treatment*, *role in the fight (witness versus non-witness), stress (pre- versus post-stress)* as factors. As post-hoc analysis, the nonparametric Mann– Whitney U test was used to compare FLX-rats with CTR-rats, and witness versus non-witness conditions. For this test, we controlled for the different amount of fights per animals by calculating the average of the frequencies and durations of behaviors of all encounters (5-7) per animal and use this average as new data points. The Wilcoxon test, on the other hand, was used to compare pre- with post-stress conditions. P<.05 (two-tailed) was considered statistically significant. Male and female data were analyzed separately.

Since relatively few litters were used, a Kruskal-Wallis test was performed to check for possible litter effects, which were not found.

## 3. Results

### 3.1 Prosocial behavior in female rats

Analysis of data revealed a trend in main effect of female rats witnessing an aggressive encounter (witness) versus when they did not (non-witness, factor *role in fight*) on time spent on active social behaviors (F_(2,39)_=2.942, p=.088), with a trend in interaction effect between the *role in fight* and the *treatment* received (F_(4,39)_=3.318, p=.07). Post-hoc analysis revealed that CTR-females that witnessed the aggressive encounters spent significantly more time on active social behaviors than when they did not witness the fights (Z=−2.43; p=.015; *d*=1.232, Figure 2A). FLX-females, on the other hand, showed no differences in time spent on active social behavior between witness and non-witness instances. While no differences were found between non-witness CTR-versus non-witness FLX-females, witness FLX-females spent significantly less time in active social behavior than witness CTR-females after an aggressive encounter (Z=−2.192; p=.030; *d*=1.474, Figure 2A).

**Figure 2.**
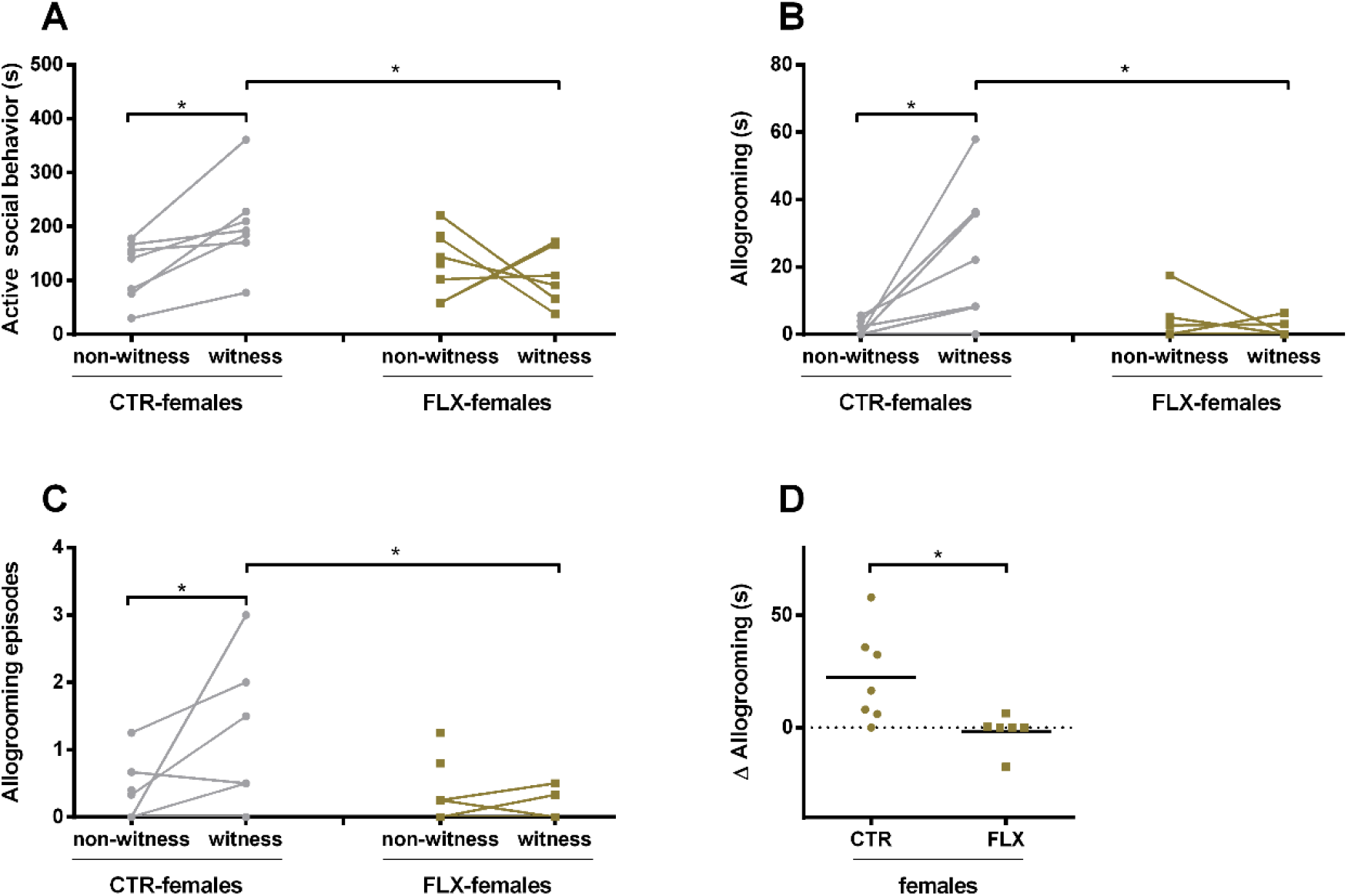
The data represents the time spent (s) on or number of episodes of a behavior by CTR- and FLX-females at adulthood in a seminatural environment (from day 1). The graphs show a comparison between instances when the CTR-rats witnessed aggressive encounters (witness) versus instances when they did not (non-witness). Data are shown as individual data points, with the lines connecting data points from the same rat calculated as the average of seconds spent on the behavior when witnessing or not-witnessing the different aggressive encounters. * p<0.05. (A) the time spent in the behavioral cluster “active social behavior”, (B) the time spent allogrooming (which is component from the behavioral cluster seen in figure 1A) (C) the number of allogroom episodes, D) the difference in time spent on allogrooming during instances as witness and as non-witness. The graph shows a comparison between CTR-females and FLX-females. (Witness CTR-females n=7, non-witness CTR-females n=8, witness FLX-females n=6, non-witness FLX-females n=8)

When the behaviors within the behavioral cluster were analyzed separately, we found a clear significant main effect of *treatment* (F_(2,39)_=5.578, p=0.027), and an interaction effect between *treatment* and *role in fight* (F_(4,39)_=11.079, p=.001) in the time spent allogrooming: witness CTR-females were allogrooming longer (Z=−2.593; p=.010; *d*=1.616, Figure 2B) and more often (trend: Z=−1.893; p=.058; *d*=1.068, Figure 2C) than non-witness CTR-females, while FLX-females that witnessed the aggressive encounters did not allogroom more or less compared to the non-witnesses. In addition, witness FLX-females spent significantly less often (Z=−2.105; p=.035; *d*=1.41, Figure 2C) and less time (Z=−2.314; p=.021; *d*=1.67, Figure 2B) in allogrooming compared to witness CTR-females. When the differences between non-witness and witness instances were calculated per rat and compared between CTR- and FLX-females, the data confirmed that CTR-females showed a significantly larger increase in allogrooming than the FLX-females that actually did not show any changes at all. (Z=−2.390; p=.017; *d*=1.671, Figure 2D). For the other behaviors within the cluster “active social behavior” (sniffing others and anogenital sniffing), no differences were found within and between CTR- and FLX-females, or witness versus non-witness (Table S1).

Also in terms of passive social behavior, there was no significant difference between witness and non-witness CTR- and/or FLX-females. However, an interaction effect of *role in fight* and *treatment* was found in the cluster “social context”, which contains all behaviors in the clusters passive social behavior and active social behavior (plus self-grooming near others) (F_(4,39)_=4.057, p=.046, Figure S1A/B): witness CTR-females showed an increase in time spent within a social context compared to non-witnesses (Z=−2.083; p=.040; *d*=1.349).

Interestingly, when the receivers of the prosocial behaviors were analyzed in more detail, it was found that although there seems to be an increase of active social behavior in CTR-females towards the losers, there was no significant difference in active social behavior towards other witnesses, non-witnesses, losers or winners in neither CTR-nor FLX-females (Figure 3).

**Figure 3.**
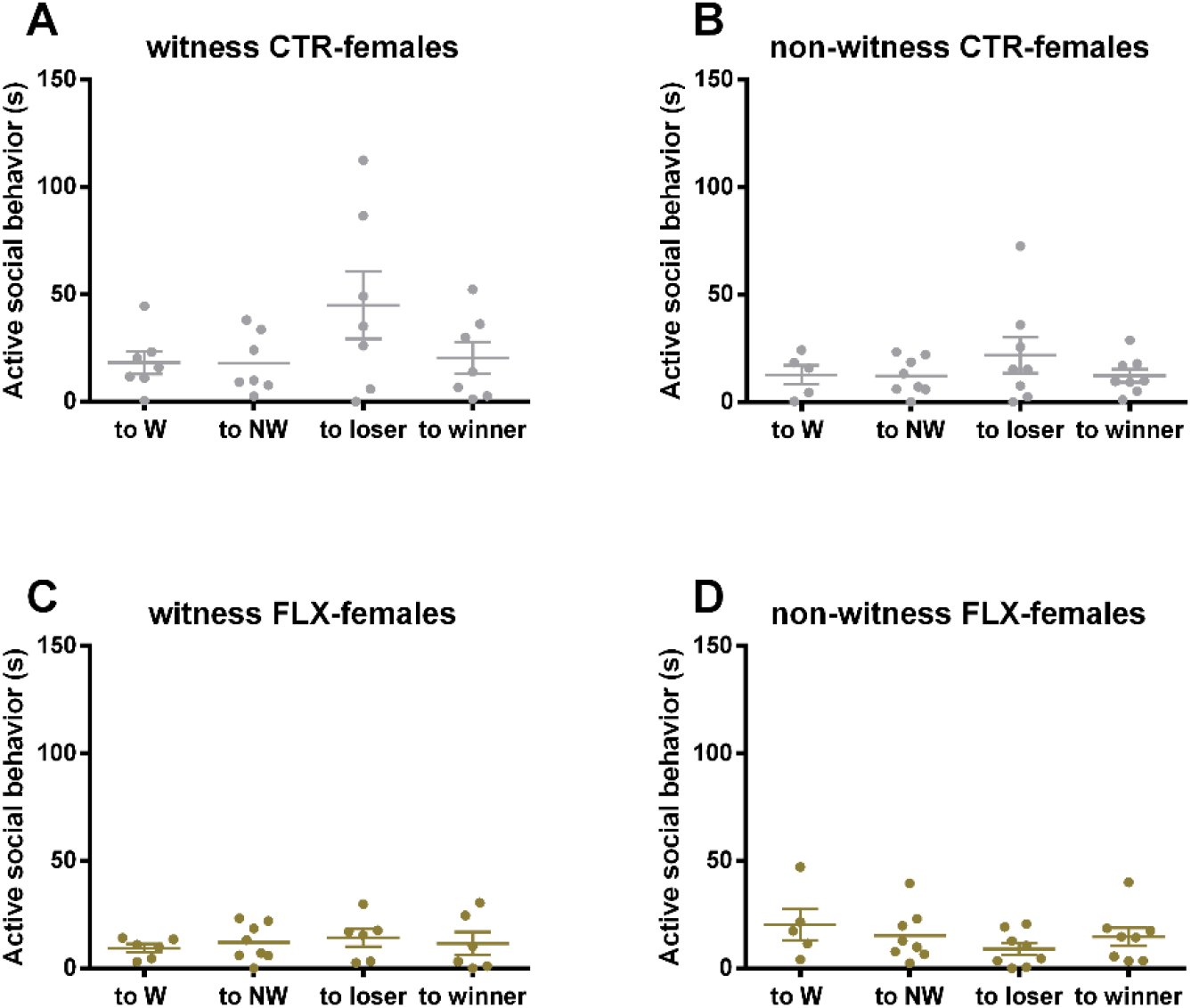
The data represents the time spent (s) on active social behavior towards witness, non-witness, losers and winners by a) witness CTR-females (n=7), B) non-witness CTR-females (n=8), C) witness FLX-females (n=6) and D) non-witness FLX-females (n=8) at adulthood in a seminatural environment (from day 1). Data are shown as individual data points (corrected by division by the number of potential receivers in that category), with the data points from the same rat calculated as the average of seconds spent on the behavior when witnessing or non- witnessing the different aggressive encounters. To W = to witness, to NW = to non-witness.

With regard to any other behavior, no differences were found between whether or not CTR- and/or FLX-females witnessed the aggressive encounter. CTR and FLX-witnesses spent, for instance, the same amount of time on general activity, passive behavior, and self-grooming as non-witnesses (Figure S1).

### 3.2 Prosocial behavior in male rats

Interestingly, analysis of data revealed that male rats do *not* show prosocial behavior after witnessing an aggressive encounter. No main effects or interaction effect of *treatment, and/or role in fight* were found on the time spent on active social behavior, nor in the separate elements within the cluster (sniffing others, allogrooming and anogenitally sniffing, Figure 4A). When the differences between non-witness and witness instances were calculated per rat and compared between CTR- and FLX-males, the data confirmed that males did not show differences in allogrooming (Figure 4B). CTR- and FLX witness and non-witness male rats also spent a similar amount of time on, for instance, passive social behavior, general activity, passive behavior, and self-grooming (Figure S2).

**Figure 4.**
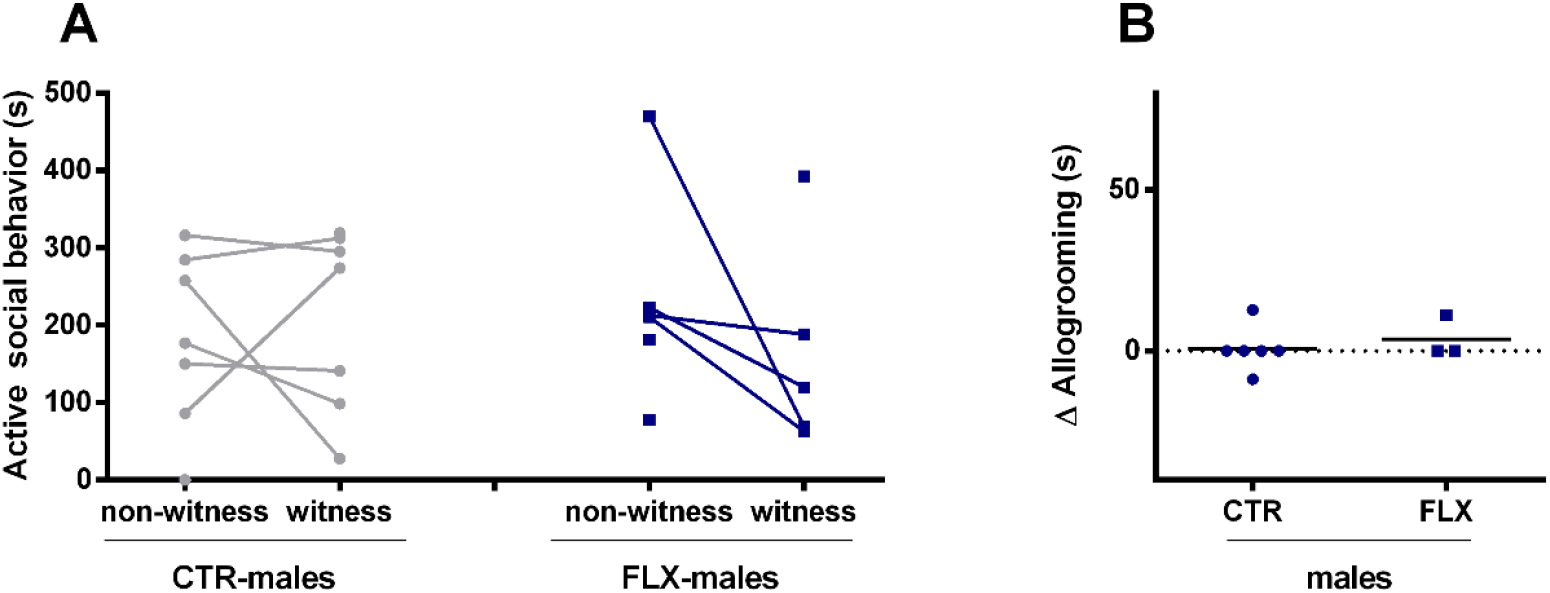
(A) The data represents the time spent (s) on active social behavior by CTR- and FLX-males at adulthood in a seminatural environment (from day 1). The graphs show a comparison between instances when the CTR-rats witnessed aggressive encounters (witness) versus instances when they did not (non-witness). Data are shown as individual data points, with the lines connecting data points from the same rat calculated as the average of seconds spent on the behavior when witnessing or not-witnessing the different aggressive encounters. (B) the difference in time spent on allogrooming (component of the cluster active social behavior) during instances as witness and as non-witness. The graph shows a comparison between CTR-males and FLX-males. (Witness CTR-males n=7, non-witness CTR-males n=7, witness FLX-males n=5, non-witness FLX-males n=6)

### 3.3 The effects of an additional stressor on prosocial behavior in female and male rats

The last aim of the study was to investigate whether an additional stressor (white-noise exposure) can alter the behaviors found in CTR- and FLX-rats. The data analysis in female rats revealed that no main effect of *stress*, or interaction effect between *stress* and other factors were found on active social behavior, or allogrooming, in CTR- and FLX- females (Figure 5A-D). Contrary to previous finding at baseline, a separate analysis of the post-stress condition, following white-noise exposure, revealed that CTR-females now did *not* show an increase in active social behavior (or the subcomponent allogrooming) when they witnessed an aggressive encounter, compared to when they did not witness the fight (Figure 5A/B). Interestingly, FLX-- females, on the other hand, even allogroomed less during instances of witnessing a fight compared to non-witnessing (Z=−2.411; p=.016; *d*=1.383, Figure 5B). Furthermore, FLX-females also no longer spent more time in a social context (Figure S3B). In fact, witness CTR-females spent significantly less time on passive social behavior compared to non-witness instances (Z=−2.641; p=.008; *d*=1.209, Figure S3A). When the behaviors in the post-stress conditions were compared to the pre-stress measures (for witness and non-witness instances separate), the data revealed that no differences between pre- and post-stress were found on any behavior in female rats.

**Figure 5.**
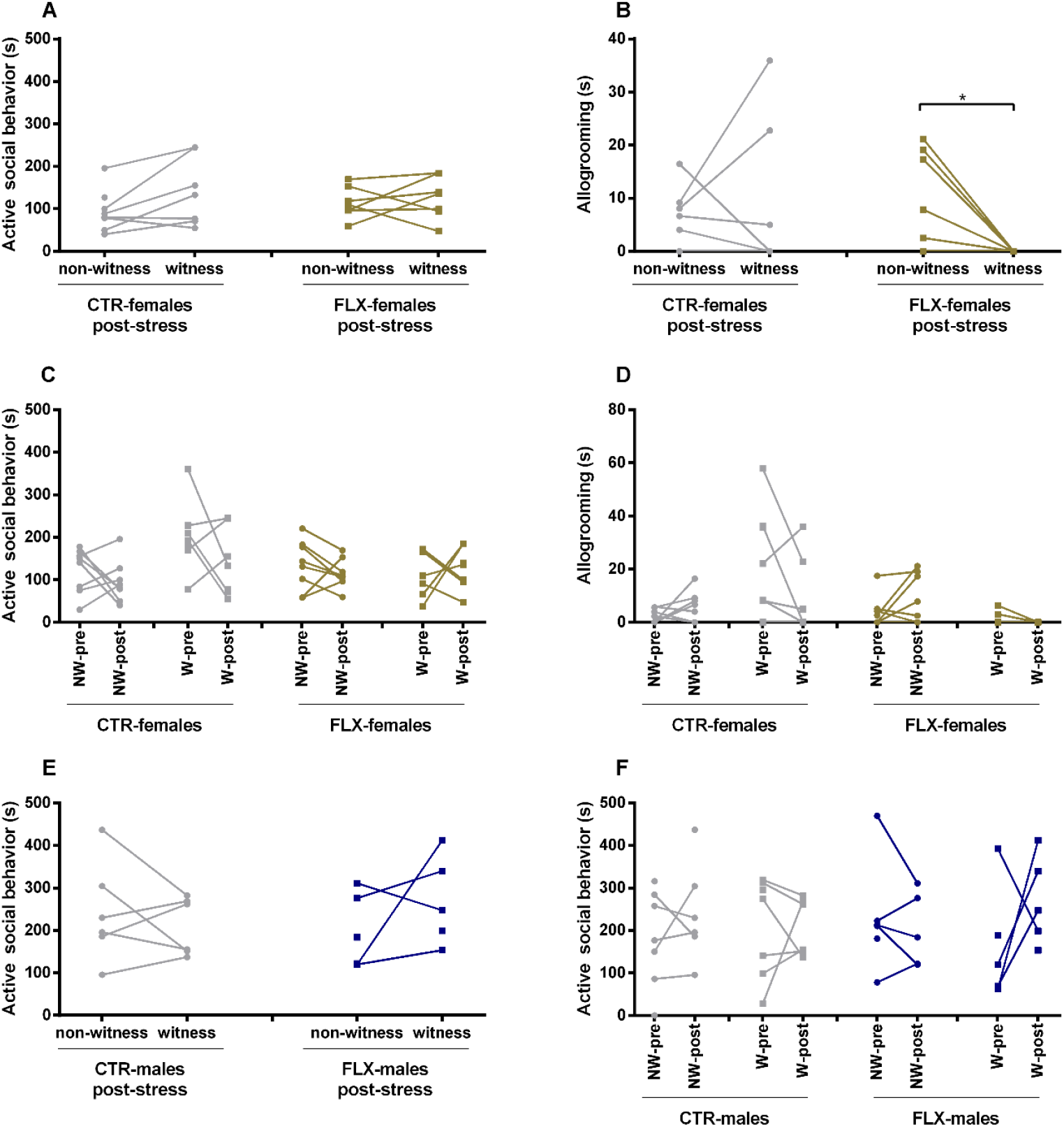
The data represents the time spent (s) on a behavior by CTR- and FLX-rats at adulthood in a seminatural environment. The graphs show a comparison between instances when the CTR-rats witnessed aggressive encounters (witness) versus instances when they did not (non-witness) (A, B, E) or between pre- versus post-stress (after white-noise exposure) conditions (C, D, F). Data are shown as individual data points, with the lines connecting data points from the same rat calculated as the average of seconds spent on the behavior when witnessing or not-witnessing the different aggressive encounters. * p<0.05. (A) the time spent in the behavioral cluster “active social behavior” of CTR- and FLX-females after white-noise exposure (post-stress) (B) the time spent allogrooming (which is component from the behavioral cluster seen in figure 5A) of CTR- and FLX-females after white-noise exposure (post-stress) (C) the time spent in the behavioral cluster “active social behavior” of CTR- and FLX-females comparing non-witness (NW) and witness (W) condition during pre- and post-stress, (D)) the time spent allogrooming of CTR- and FLX-females comparing non-witness (NW) and witness (W) condition during pre- and post-stress (E) the time spent in the behavioral cluster “active social behavior” of CTR- and FLX-males after white-noise exposure (post-stress) (F) the time spent in the behavioral cluster “active social behavior” of CTR- and FLX-males comparing non-witness (NW) and witness (W) condition during pre- and post-stress, (Pre-stress: Witness CTR-males n=7, non-witness CTR-males n=7, witness FLX-males n=5, non-witness FLX-males n=6, witness CTR-females n=7, non-witness CTR-females n=8, witness FLX-females n=6, non-witness FLX-females n=8, post-stress: Witness CTR-males n=6, non-witness CTR-males n=6, witness FLX-males n=5, non-witness FLX-males n=6, witness CTR-females n=7, non-witness CTR-females n=8, witness FLX-females n=7, non-witness FLX-females n=8)

In addition, no main effects of *stress*, or interaction effects between *stress, treatment,* and/or *role in fight* were found on any other behavior in female rats (Figure S3 and S4), except for an interaction effect between *stress* and *role in fight* on general activity (F_(2,39)_=5.637, p=0.019), probably caused by an increase in general activity of witness FLX-females compared to non-witness FLX-females in the post-stress condition (Z=−2.315; p=.021; *d*=1.586, not significantly different from CTR-females, Figure S3C). Also for the time spent and number of episodes of conflict behaviors, a main effect on *stress* (duration: F_(2,39)_= 19.442, p>0.001, episodes: F_(2,39)_=13.501, p<0.001), and *stress*-*role in fight* (duration: F_(4,39)_=4.947, p=0.027, episodes: F_(2,39)_=5.957, p=0.016) was found, but post-hoc analysis did not show any differences.

In male rats, on the other hand, a similar mixed models analysis revealed that there was a significant main effect of *stress* (F_(2,39)_= 5.188, p=0.025), and an interaction effect of *stress* and *role in fight* (F_(4,39)_= 4.363, p=0.04), and on *stress*, *treatment*, and *role in fight* (F_(8,39)_= 4.780, p= 0.032) on time spent on active social behaviors (Figure 5E/F). Post-hoc analysis, however, did not show any significant differences between non-witness versus witness, or pre-versus post-stress condition in CTR- and/or FLX-male rats on active social behavior (Figure 5E/F). Also on the time spent in a social context, a significant main effect was found on *stress* (F_(2,39)_=5.059, p=0.27), and interaction effects between *stress, role in fight* and *treatment* (F_(8,39)_=5.923, p=0.017) were found (Figure S5 and S6). Again, post-hoc analysis did not reveal any particular differences.

Only in terms of conflict behavior, a main effect on *role in fight* (F_(2,39)_=4.213, p=0.043), and an interaction effect of *stress* and *role in fight* (F_(4,39)_=7.365, p=0.008) was found. Post-hoc analysis revealed that non-witness FLX-males spent significantly more time in conflict behaviors than non-witness CTR-males after white-noise exposure, but this effect was clearly caused by an artifact (figure S5F).

No differences were found on any of the other behaviors, except for a main effect of *stress* on the time spent on passive behavior (F_(2,39)_=4.353, p=0.040), an effect that was most likely caused by a decrease in passive behavior in FLX-males compared to CTR-males (Z=− 2.143; p=.032; *d*=1.2) after white-noise exposure, and compared to FLX-males in the pre-stress condition (Z=−1.963; p=.050; *d*=1.3, Table S1). In addition, a main effect of *stress* was found on episodes of self-grooming (F_(2,39)_=6.841, p=0.011), but this effect was no longer confirmed by post-hoc analysis.

## 4. Discussion

The first aim of the study was to explore if rats are capable of showing prosocial behavior. As mentioned before, prosocial behavior is defined as behavior that is intended to benefit another conspecific, and includes helping and consolation behavior. In our study, we investigated the possibility that rats show consolation behavior, which should be visible as an increase in affiliative contact directed towards presumed distressed conspecifics in response to an aggressive encounter that took place in close vicinity. Therefore, we compared the behavior of rats that witnessed versus those that did not witness the aggressive encounters. It was hypothesized that witnesses, in contrast to non-witnesses, were able to observe the consequences of a stressful event that potentially triggered distress in conspecifics (both those involved in the fights and third-party bystanders).

Our results indicate that female rats do indeed show post-conflict prosocial behavior. CTR-females, but not CTR-males, showed higher levels of social behavior towards conspecifics after instances when they had witnessed an aggressive encounter compared to after instances when they had not witnessed the fight. This suggests that witness CTR-females seek social interaction after a stressful social conflict situation, and thereby most likely either try to seek comfort from, or offer social relief to their conspecifics which experienced discomfort caused by a social conflict in the environment. Previous studies in other rodent species have suggested that consolation behavior, a form of prosocial behavior, is mostly expressed as allogrooming behavior [46, 50]. In addition to a general effect in social context and active social behavior, our results also showed an increase in allogrooming behavior after witnessing a fight. This suggests that the increase in active social behavior, including allogrooming, might have offered social relief to the conspecifics in distress, and could thus be considered post-conflict prosocial behavior.

The second aim of the study was to investigate whether perinatal SSRI exposure can affect the expression of prosocial behavior in rats at an adult age, which could be an indicator for deficits in social behavior. We found that FLX-females, in contrast to CTR-females, did not show any signs of post-conflict prosocial behavior in our experiment in the seminatural environment. This supports our evidence that perinatal fluoxetine exposure does affect prosocial behavior in female rats. Though, we can conclude that males and females cope differently with stressors, because CTR-males did not show any signs of post-conflict prosocial behavior. It is therefore impossible to draw conclusions about whether or not perinatal fluoxetine exposure also affects male prosocial behavior.

The third aim of the study was to see if an additional stressor would alter the occurrence of prosocial behavior. Indeed, the increase in active social behavior in CTR-females that occurred following an aggressive encounter (indicated as post-conflict prosocial behavior) diminished after a stressful white-noise exposure. Additionally, in response to the combined stressors of white-noise *and* an aggressive encounter, no differences in behavior were found for CTR- and FLX-male rats.

Collectively, these results suggest that the perinatal SSRI exposure has sex- and context-dependent effects on post-conflict prosocial behavior.

### 4.1 Consolation behavior

Although our results indicate the existence of prosocial behavior after instances of witnessing aggressive encounters, at least in females, it is unclear whether this prosocial behavior can be called *consolation* behavior. In order to be defined *consolation* behavior, the prosocial behavior should have a stress-lowering effect in the conspecifics [46, 51]. Since we were not able to measure corticosterone levels, nor physiological parameters before and after the aggressive encounters or after the received prosocial behavior, we cannot be absolutely certain that the observed behavior is indeed *consolation* behavior. However, the fact that the prosocial behavior is solely seen after instances of *witnessing* an aggressive encounter is in line with studies on consolation behavior by third-party bystanders in different species. In bonobos, for instance, more consolation behavior was shown to be displayed when an animal was closer to a conflict [41]. In addition, the anti-stress effect of allogrooming has previously also been shown to reduce heart rates in several species [52–54]. Besides, allogrooming is not a typical response to first-party stress, and therefore would not be expected as a pure consequence of contagious stress, anxiety or fear. Altogether, this suggests that the increase in active social behavior, including allogrooming, found in our study might offer social relief to their conspecifics, and could be considered consolation behavior.

In our study, the displayed post-conflict prosocial behavior was mostly directed at individuals that were themselves not involved in the aggressive encounter (third-party bystanders, or ‘others’). CTR-females did not show significantly more active social behaviors towards the others (witness or non-witness) than towards the losers and winners. Initially, we expected that consolation behavior would take place more towards the losers, as also shown in other the species [41, 42]. However, it is not unthinkable that prosocial behavior is *also* aimed at the other rats which probably suffered from distress caused by the social conflict within their environment. The conflict could spread a change in general arousal of rats through the environment, which could then explain why consolation is offered to all rats, instead of solely towards the losers A similar finding was presented for Asian elephants that offered consolation to other bystanders [44]. In addition, rats that lose the fight might actually prefer to withdraw from social contact after defeat and are therefore not actively pursued. In fact, it has been shown that after a serious social defeat, rats lower their general activity and refrain from social contact, even with non-aggressive conspecifics [55, 56].

Another explanation could be that the spontaneous aggressive encounters in our setting were not severe enough to evoke consolation behavior towards the losers. None of the rats involved in the aggressive encounters in our study suffered visible physical damage nor seemed to change their behavior remarkably. Our study was an observational study in a natural setting, meaning that we did not have control over the timing or severity of the aggressive encounters. This could be the reason why our effects are limited in variety of behaviors, and why we did not observe post-conflict prosocial behavior in CTR-males.

At last, it should be mentioned that the increase in active social behavior could not be explained by the increased arousal in female rats after a stressful event, as shown by the fact that other general activity measures such as running/walking and non-social exploration were not affected in these animals.

### 4.2 How does perinatal fluoxetine exposure affect prosocial behavior?

It is evident in our FLX-females that perinatal SSRI exposure affects prosocial behavior. The response to witnessing aggressive encounters seen in CTR females, is absent in almost all of the FLX-exposed females. Behavior of FLX-females seems unaltered by instances of witnessing an aggressive encounter, whereas CTR-females react with increased active social behavior, including allogrooming. This indicates possible deficiencies in the prosocial response of FLX-females. The question remains why FLX-females (and FLX-males) lack the “normal” response to witnessing an aggressive encounter. One hypothesis is that perinatal SSRI exposure alters the responses to stressors via changing control mechanisms involved in the regulation of the negative feedback of the hypothalamic-pituitary-adrenal (HPA) axis. Corticosterone and other glucocorticoids play an important role in regulating HPA axis functionality. One study showed that prenatal exposure to fluoxetine can increase corticosterone responses to acute and continuous stressors and at the same time induce a state of glucocorticoid resistance in adult female mice [57]. Interestingly, another study found that a social interaction with a novel conspecific was not sufficient to induce higher levels of corticosterone in perinatally fluoxetine exposed rats [30], but they still found increased levels of corticosteroid-binding globulin [30, 58]. Together, this suggests that the HPA response is indeed modified after perinatal SSRI exposure. Although not investigated in our study, this altered HPA response to stress could underlie the differences between CTR- and FLX-females found in our study on prosocial behavior.

In fact, Ben-Ami Bartal et al. found an inverted U-shape effect of stress and another form of prosocial behavior, helping behavior. Both low levels of negative arousal (by midazolam treatment, a benzodiazepine anxiolytic acting on the HPA axis) *and* high corticosterone responses resulted in impairment of helping behavior in rats [59]. As mentioned above, our aggressive encounters might not have been intense enough to trigger a strong stressful response, and thereby induce prosocial behavior in the CTR-male rats. That high levels of stress impair post-conflict prosocial behavior, might to the contrary be an explanation for the lack of prosocial behavior in FLX-rats which have increased HPA responses to stress. In line with another study showing that prosocial behavior was negatively affected by HPA activity [60], Ben-Ami Bartal et al. concluded that the HPA reactivity results in antagonistic effects on helping behavior [59]. They might just get “too afraid to help” [61]. In fact, individuals with the short allele polymorphism of the serotonin transporter gene regulatory region (5-HTTLPR) also have higher HPA reactivity [62] *and* lower prosocial tendencies [61]. Again, this suggests a clear relationship between serotonin, the HPA axis and prosocial behavior. In the case of our study, the stress caused by witnessing an aggressive encounter could be severe enough to evoke a different HPA reactivity in FLX-females compared to CTR-females. This could result in the impairment of prosocial behavior.

### 4.3 An additional stressor and prosocial behavior

The hypothesis that high levels of stress impairing prosocial behavior might also explain the lack of post-conflict prosocial behavior found in CTR-females after the exposure to white-noise. While witnessing an aggressive encounter at baseline results in prosocial behavior because it caused only moderate levels of stress in the third-party bystanders, the exposure to white-noise could have induced higher levels of stress which subsequently block the display of post-conflict prosocial behavior in the post-stress condition. In the case of FLX-females, the additional stressor even impaired the post-conflict prosocial behavior more than before the white-noise. This finding was in line with our previous findings in terms of social behavior in which FLX-females switched from resting socially to more solitary after a stressful event [31]. In the current study, this effect was also seen in CTR-females and CTR-males.

Previously, we have found that the exposure to white-noise induces self-grooming behavior in FLX-males which was hypothesized to be due to alterations in stress-coping behavior [31]. In the current study, we do not observe elevated levels of self-grooming in CTR- and FLX-males after witnessing aggressive encounters in the pre- and post-stress conditions. An explanation can be found in that the social stressor in the current experiment could have been too light and short lasting to induce self-grooming. In addition, the behavior was observed immediately after the white-noise exposure in the previous study, while in the current study we observed the behavior after the aggressive encounters, which occurred later on the day.

Unpublished data revealed that the increased self-grooming behavior of FLX-males slowly attenuated over time (measured up to 3 hours after the white-noise), with a large variability between individuals. Thus, we conclude that an additional stressor did not affect the behavior of CTR- and FLX-rats in general after aggressive encounters. However, it did affect post-conflict prosocial behavior of CTR-females, which is in line with the hypothesis that an increased HPA reactivity impairs the motivation to show prosocial behavior.

### 4.4. Strengths and limitation of the study

Since our study uses an unconventional design, we would like to point out some strengths and limitations of the study. Due to the complex nature of the experimental design, our study has a rather small sample size comprised of offspring from only 6 litters. It would be challenging to include offspring from separate dams in one study. Instead, we have taken all measures possible to limit the use of litters within one cohort. The advantage of this design is that we were able to investigate and confirm that there was no effect of litter on the behavioral outcomes. The unconventional design, however, also results in the major strength of our study: the use of a seminatural setting that allows for the observation of natural spontaneously occurring behavior.

The rats are able to express their full repertoire of natural behaviors, or to *not* express certain behaviors. Not only does this method allow to observe the real natural response of an animal, it also opens the possibility to study different behaviors in relation to each other in a certain context.

## 5. Conclusion

In summary, we conclude that female rats are able to show prosocial behavior in response to witnessing aggressive encounters between conspecifics in a seminatural environment. We suggest that this post-conflict prosocial behavior is possibly consolation behavior, but more research would be needed to confirm this. Male rats, on the other hand, did not show post-conflict prosocial behavior in our test setting, suggesting that either they do not show consolation behavior, or our test set-up is not sufficient to detect this behavior in males. Our observational study was the first to show this behavior in rats in a natural setting with spontaneous occurring aggressive encounters.

In addition, we showed that perinatal fluoxetine exposure impairs the display of post-conflict prosocial behavior in female rats. We hypothesize that this is caused by an increased HPA reactivity, since an additional stressor (exposure to white-noise) also disrupts the post-conflict prosocial behavior seen in CTR-females. Further research in the HPA reactivity is necessary to confirm this hypothesis.

The effects of perinatal fluoxetine exposure on prosocial behavior could provide additional evidence that SSRI treatment during pregnancy could contribute to the risk for social impairments in the offspring. In conclusion, the experimental set-up used in this study may be of great help for the study of psychiatric disorders, in particular to the aspect of prosocial behavior.

## 6. Acknowledgments

Financial support was received from Helse Nord #PFP1295-16, Norway. DJH was supported by the KNAW ter Meulen travel grant, the Netherlands.

We also would like to thank Patty Huijgens for the rat images, and Aslaug Angelsen for the unpublished data on self-grooming. In addition, we thank Ragnhild Osnes, Carina Sørensen, Nina Løvhaug, Katrine Harjo, and Remi Osnes for their excellent care of the animals.

