## Supplemental materials for "Third-party prosocial behavior in adult female rats is impaired after perinatal fluoxetine exposure"

[3] Regional Health Authority of North Norway

### Table S2: Information dams and offspring

Copy from Houwing et al. 2019

#### A: Treatment dams and cage distribution offspring

| **Dams** | Treatment | Number offspring | Male-Cage 1 | Male-Cage 2 | Female-Cage 3 | Female-Cage 4 | Female-Cage 5 | Female-Cage 6 |
|---|---|---|---|---|---|---|---|---|
| F1 | FLX | 7 | OM1<br>OM2<br>OM3 | | OF2<br>OF3 | OF1<br>OF4 | | |
| F2 | FLX | 9 | OM5<br>OM6<br>OM7 | OM4<br>OM8 | OF5<br>OF8 | OF6<br>OF7 | | |
| F3 | CTR | 13 | OM9<br>OM12 | OM10<br>OM11 | OF12<br>OF16 | OF9<br>OF15 | OF11<br>OF14 | OF10<br>OF13<br>OF17 |
| F4 | CTR | 6 | OM13<br>OM14<br>OM15 | | OF18<br>OF19<br>OF20 | | | |
| F5 | FLX | none | | | | | | |
| F6 | FLX | none | | | | | | |
| F7 | CTR | none | | | | | | |
| F8 | CTR | 15 | OM16<br>OM17 | OM20<br>OM21<br>OM25 | OF21<br>OF23 | OF22<br>OF24<br>OF25 | | |
| F9 | FLX | 8 | OM29<br>OM30<br>OM31 | | OF27<br>OF29 | OF26<br>OF28<br>OF30 | | |
| F10 | FLX | dead | | | | | | |

*Abbreviations: M = male, F = female, CTR = methylcellulose, FLX = fluoxetine, OM = male offpsring, OF = female offspring*

#### B: Experimental design seminatural environment

| **Colony** | **M 1** | **M 2** | **M 3** | **M 4** | **F 1** | **F 2** | **F 3** | **F 4** |
|---|---|---|---|---|---|---|---|---|
| SNE1 | OM1 | OM5 | OM9 | OM13 | OF2 | OF5 | OF12 | OF21 |
| SNE2 | OM2 | OM6 | OM12 | OM16 | OF3 | OF8 | OF13 | OF22 |
| SNE3 | OM3 | OM7 | OM15 | OM17 | OF1 | OF6 | OF15 | OF24 |
| SNE4 | OM8 | OM10 | OM18 | OM30 | OF7 | OF14 | OF20 | OF26 |

*Abbreviations: SNE= seminatural environment cohort, M = male, F = female, OM = male offpsring, OF = female offspring*

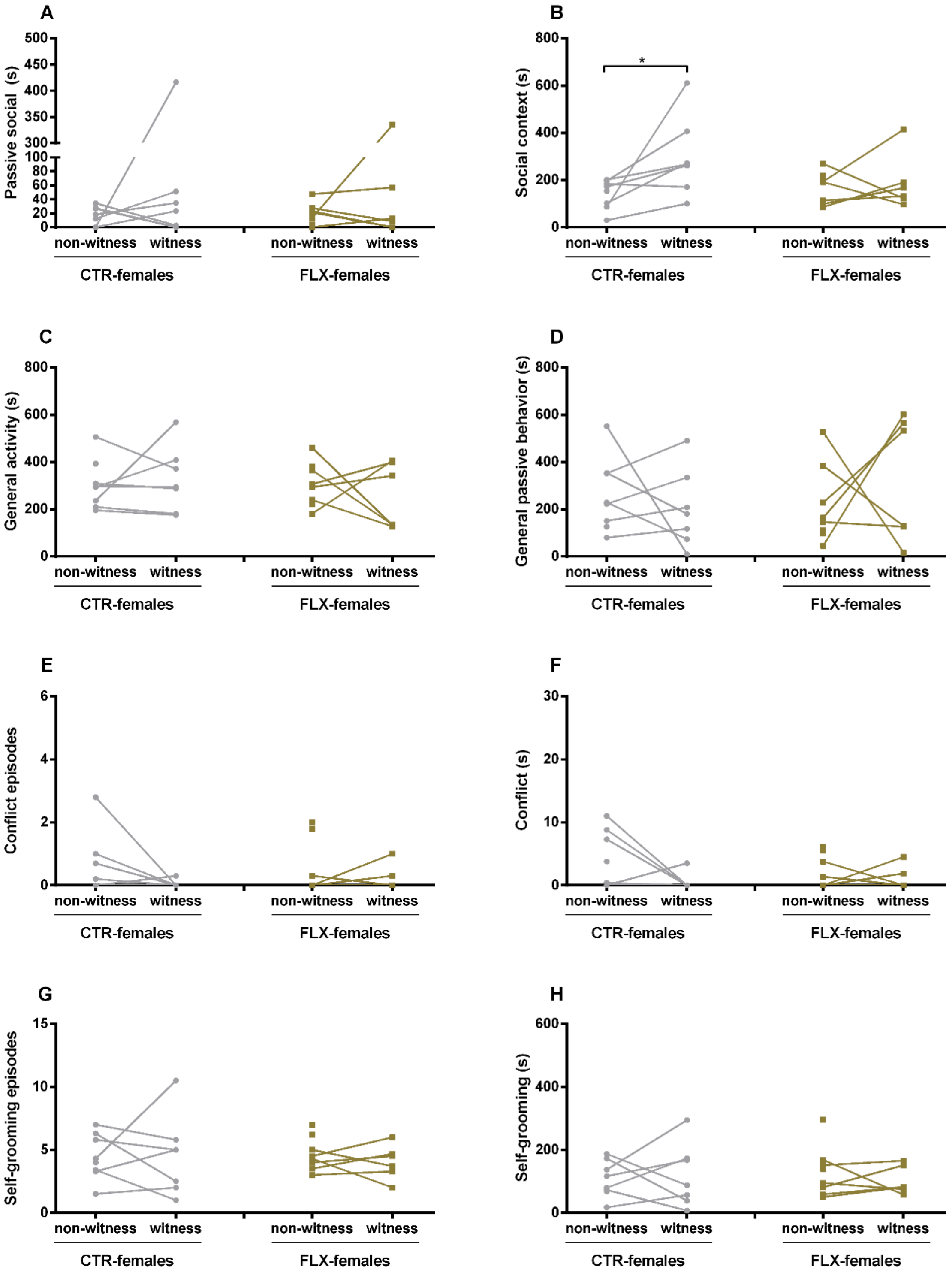


*Figure S1. The data represents either the time spent (s) on a behavior or the number of episodes of the behavior performed by CTR- and FLX-female rats at adulthood in a seminatural environment (from day 1).. The graphs show a comparison between instances when the rats witnessed aggressive encounters (witness) versus instances when they did not (non-witness). ). Data are shown as individual data points, with the lines connecting data points from the same rat calculated as the average of seconds spent on the behavior when witnessing or not-witnessing the different aggressive encounters. * p<0.05.*

*(A) time spent on passive social behavior, (B) time spent in a social context, (C) time spent being generally active, (D) time spent being generally passive, (E) number of conflict episodes, (F) time spent in conflict situations, (G) number of self-groom episodes, and (H) time spent self-grooming.*

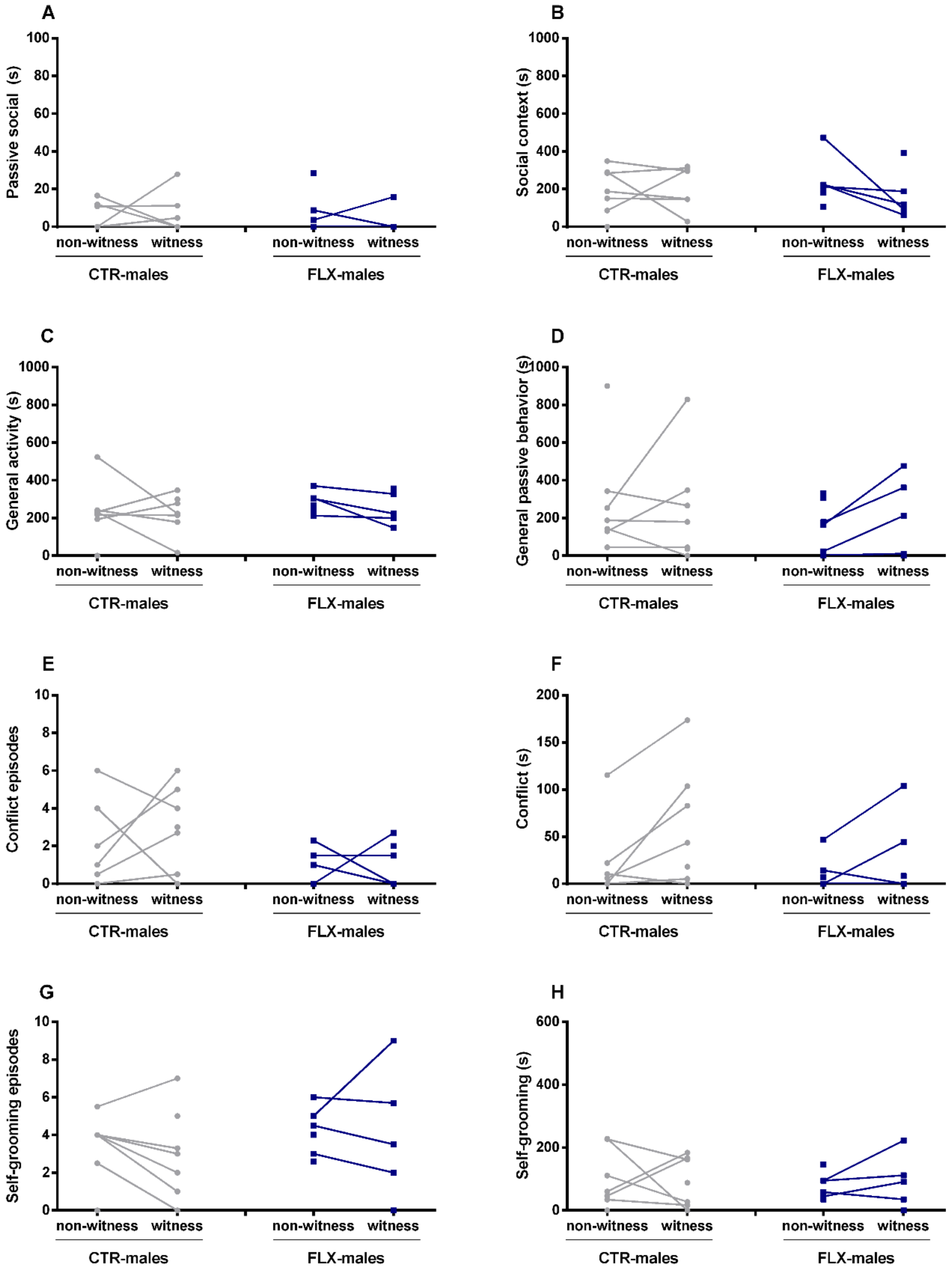


*Figure S2. The data represents either the time spent (s) on a behavior or the number of episodes of the behavior performed by CTR- and FLX-male rats at adulthood in a seminatural environment (from day 1).. The graphs show a comparison between instances when the rats witnessed aggressive encounters (witness) versus instances when they did not (non-witness). ). Data are shown as individual data points, with the lines connecting data points from the same rat calculated as the average of seconds spent on the behavior when witnessing or not-witnessing the different aggressive encounters.*

*(A) time spent on passive social behavior, (B) time spent in a social context, (C) time spent being generally active, (D) time spent being generally passive, (E) number of conflict episodes, (F) time spent in conflict situations, (G) number of self-groom episodes, and (H) time spent self-grooming.*

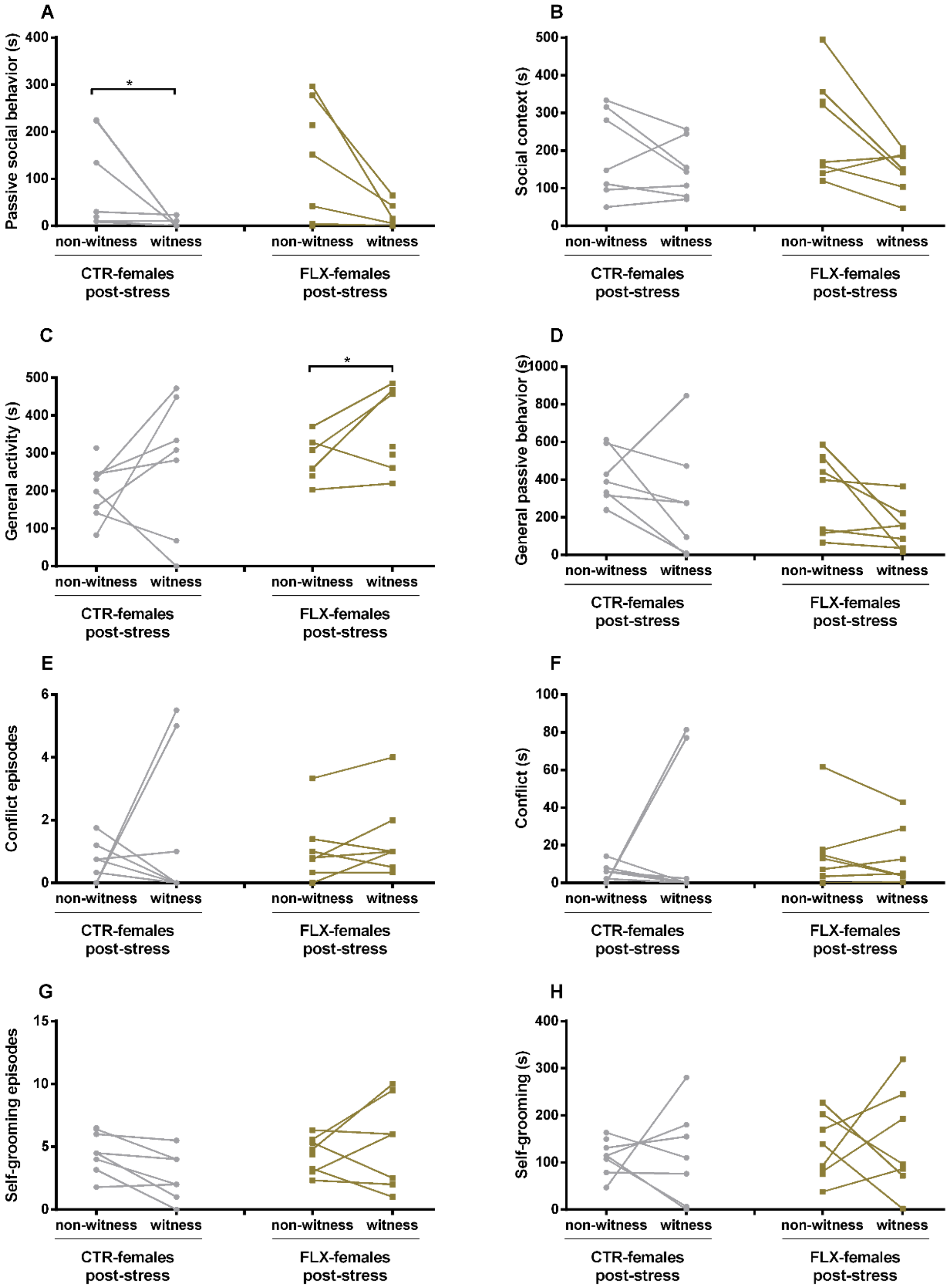


*Figure S3. The data represents either the time spent (s) on a behavior or the number of episodes of the behavior performed by CTR- and FLX-female rats at adulthood in a seminatural environment after white-noise exposure (post-stress). The graphs show a comparison between instances when the rats witnessed aggressive encounters (witness) versus instances when they did not (non-witness). Data are shown as individual data points, with the lines connecting data points from the same rat calculated as the average of seconds spent on the behavior when witnessing or not-witnessing the different aggressive encounters. * $p<0.05$.*

*(A) time spent on passive social behavior, (B) time spent in a social context, (C) time spent being generally active, (D) time spent being generally passive, (E) number of conflict episodes, (F) time spent in conflict situations, (G) number of self-groom episodes, and (H) time spent self-grooming.*

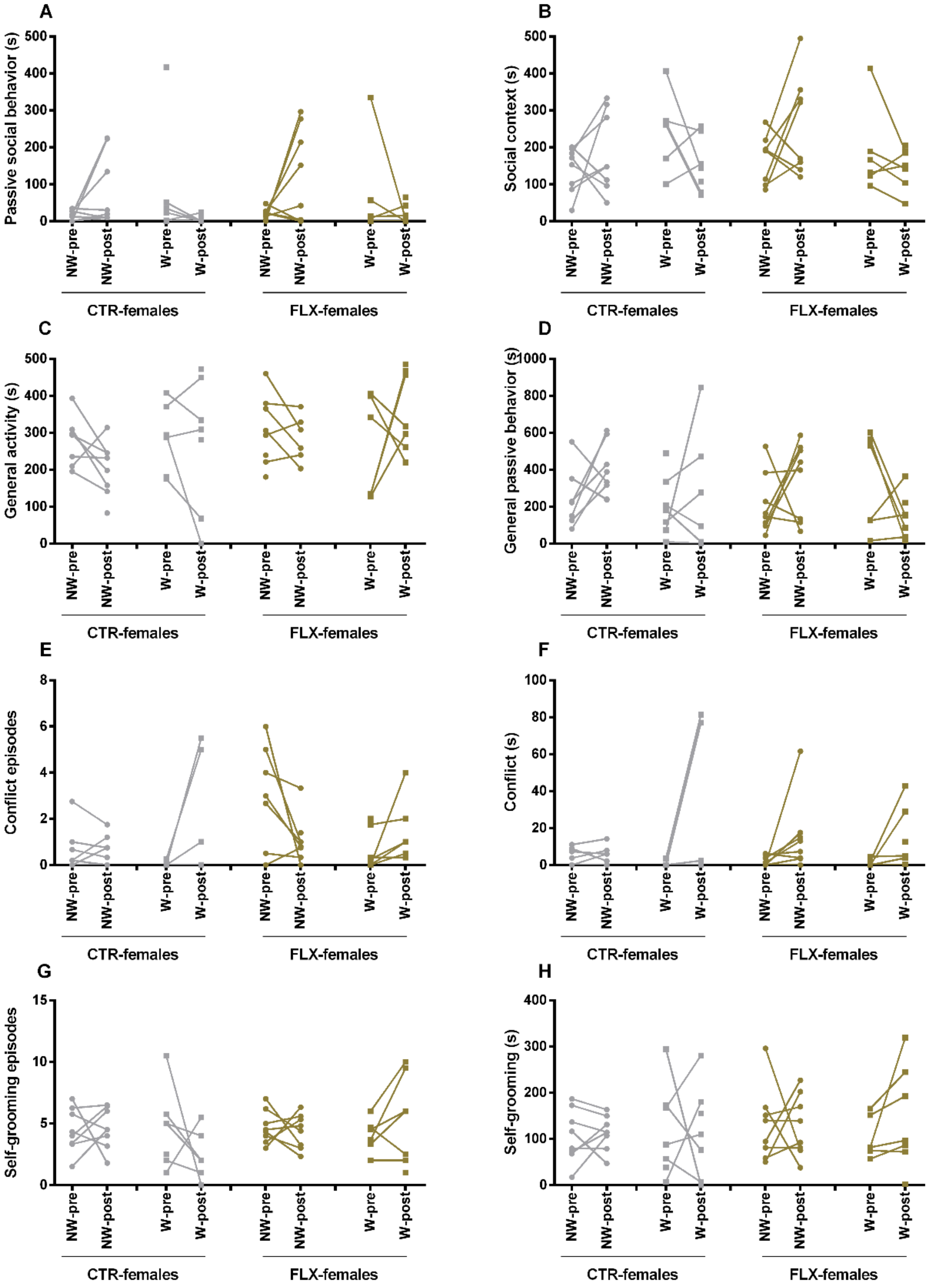


*Figure S4. The data represents either the time spent (s) on a behavior or the number of episodes of the behavior performed by CTR- and FLX-female rats at adulthood in a seminatural environment before (pre) and after (post) white-noise exposure. The graphs show a comparison between pre and post-stress conditions for instances when the rats witnessed aggressive encounters (witness, W) and instances when they did not (non-witness, NW). Data are shown as individual data points, with the lines connecting data points from the same rat calculated as the average of seconds spent on the behavior when witnessing or not-witnessing the different aggressive encounters.*

*(A) time spent on passive social behavior, (B) time spent in a social context, (C) time spent being generally active, (D) time spent being generally passive, (E) number of conflict episodes, (F) time spent in conflict situations, (G) number of self-groom episodes, and (H) time spent self-grooming.*

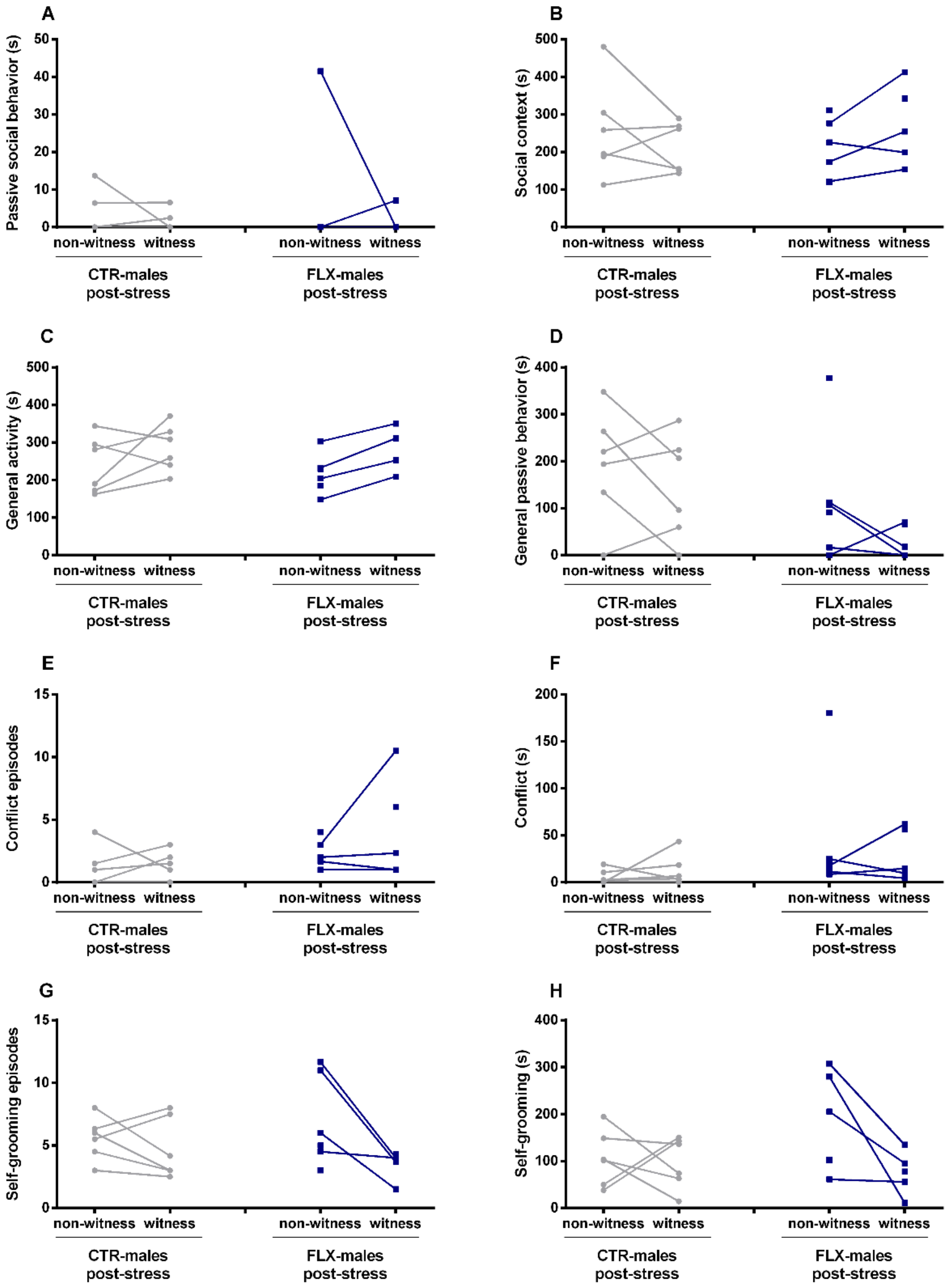


*Figure S5. The data represents either the time spent (s) on a behavior or the number of episodes of the behavior performed by CTR- and FLX-male rats at adulthood in a seminatural environment after white-noise exposure (post-stress). The graphs show a comparison between instances when the rats witnessed aggressive encounters (witness) versus instances when they did not (non-witness). Data are shown as individual data points, with the lines connecting data points from the same rat calculated as the average of seconds spent on the behavior when witnessing or not-witnessing the different aggressive encounters.*
*(A) time spent on passive social behavior, (B) time spent in a social context, (C) time spent being generally active, (D) time spent being generally passive, (E) number of conflict episodes, (F) time spent in conflict situations, (G) number of self-groom episodes, and (H) time spent self-grooming.*

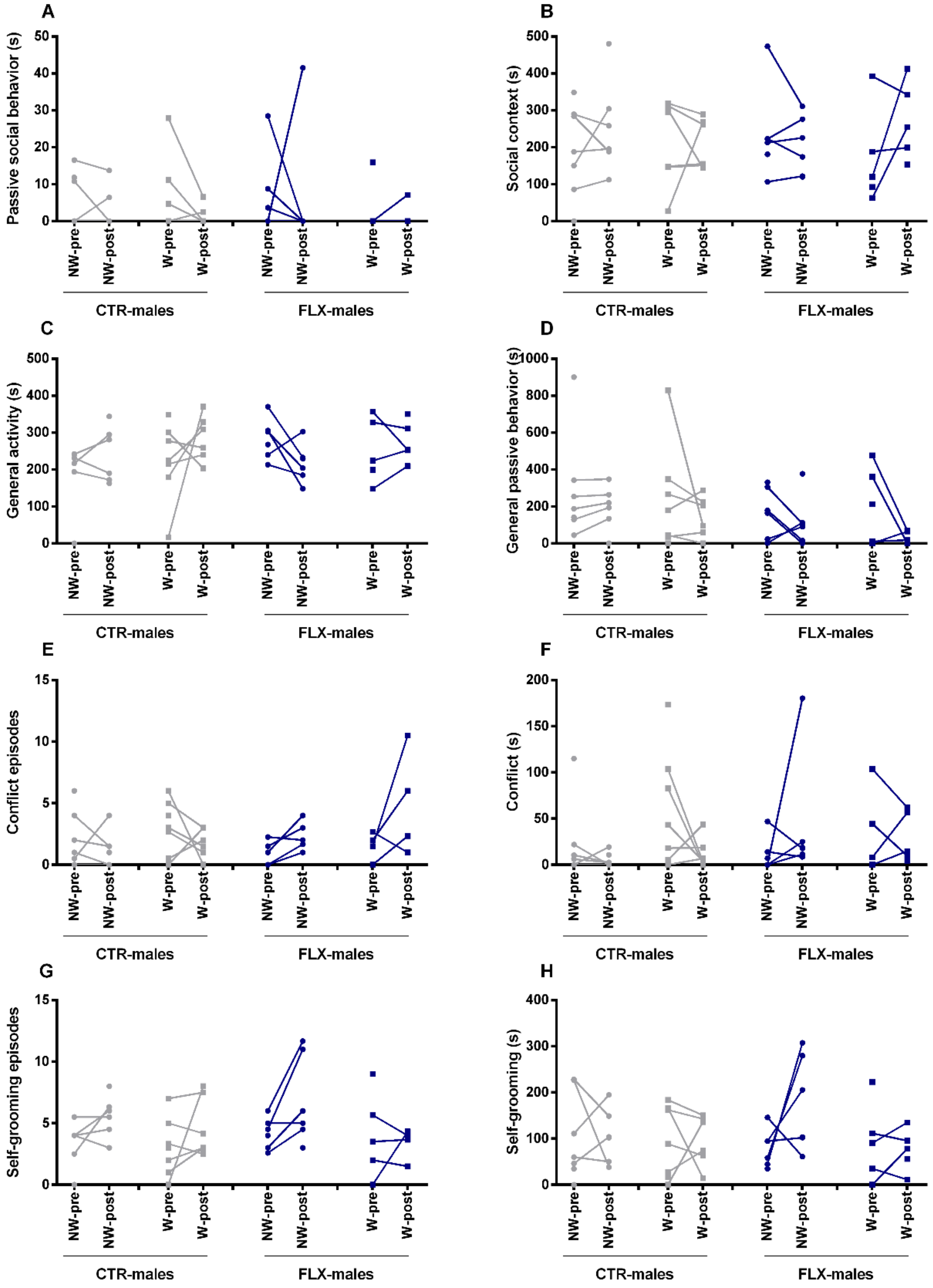


*Figure S6. The data represents either the time spent (s) on a behavior or the number of episodes of the behavior performed by CTR- and FLX-male rats at adulthood in a seminatural environment before (pre) and after (post) white-noise exposure. The graphs show a comparison between pre and post-stress conditions for instances when the rats witnessed aggressive encounters (witness, W) and instances when they did not (non-witness, NW). Data are shown as individual data points, with the lines connecting data points from the same rat calculated as the average of seconds spent on the behavior when witnessing or not-witnessing the different aggressive encounters.*

*(A) time spent on passive social behavior, (B) time spent in a social context, (C) time spent being generally active, (D) time spent being generally passive, (E) number of conflict episodes, (F) time spent in conflict situations, (G) number of self-groom episodes, and (H) time spent self-grooming.*
