## Supplementary Table S1 for "Third-party prosocial behavior in adult female rats is impaired after perinatal fluoxetine exposure"

The data represents average time or average number of episodes performing named behaviour +/-SEM.

Differences compared to CTR-rats are highlighted where Mann Whitney U p<0.05

Differences compared to non-witnesses are highlighted where Mann Whitney U p<0.05

| | Baseline | | | | | | | | Post-stress | | | | | | | |
|---|---|---|---|---|---|---|---|---|---|---|---|---|---|---|---|---|
| | Female | | | | Male | | | | Female | | | | Male | | | |
| | CTR | | FLX | | CTR | | FLX | | CTR | | FLX | | CTR | | FLX | |
| | Non-Witness | Witness | Non-Witness | Witness | Non-Witness | Witness | Non-Witness | Witness | Non-Witness | Witness | Non-Witness | Witness | Non-Witness | Witness | Non-Witness | Witness |
| n | 8 | 7 | 8 | 6 | 7 | 7 | 6 | 5 | 8 | 7 | 8 | 7 | 6 | 6 | 6 | 5 |
| Time in | | | | | | | | | | | | | | | | |
| **Active social behavior** | **122.6** ± 18.7 | **203.4** ± 32.0 | **134.2** ± 20.9 | **107.1** ± 21.9 | **181.6** ± 43.0 | **209.7** ± 44.8 | **229.1** ± 52.8 | **166.4** ± 60.8 | **95.0** ± 17.3 | **140.0** ± 30.1 | **113.9** ± 12.1 | **126.4** ± 18.9 | **241.5** ± 47.9 | **209.6** ± 27.7 | **188.7** ± 34.9 | **270.4** ± 47.0 |
| **Passive Social behavior** | **15.1** ± 4.9 | **75.5** ± 57.3 | **20.1** ± 5.2 | **68.9** ± 53.9 | **5.6** ± 2.7 | **6.3** ± 3.9 | **6.8** ± 4.6 | **3.2** ± 3.2 | **82.6** ± 34.2 | **4.9** ± 3.5 | **123.2** ± 45.1 | **18.4** ± 9.7 | **3.4** ± 2.3 | **1.5** ± 1.1 | **6.9** ± 6.9 | **1.4** ± 1.4 |
| **Social context** | **140.4** ± 21.7 | **298.1** ± 63.2 | **170.5** ± 22.8 | **186.7** ± 47.4 | **192.3** ± 46.9 | **221.4** ± 43.2 | **236.6** ± 50.6 | **171.1** ± 59.0 | **185.6** ± 38.4 | **150.9** ± 28.3 | **261.4** ± 47.3 | **146.4** ± 21.0 | **256.6** ± 52.1 | **211.8** ± 27.8 | **204.6** ± 32.6 | **272.3** ± 47.1 |
| **General active behavior** | **304.9** ± 36.4 | **326.4** ± 52.1 | **305.7** ± 32.8 | **256.8** ± 56.9 | **234.3** ± 57.8 | **223.0** ± 40.6 | **283.1** ± 22.7 | **251.2** ± 39.4 | **202.0** ± 25.7 | **273.3** ± 67.7 | **208.5** ± 32.7 | **358.0** ± 41.4 | **240.8** ± 30.9 | **285.0** ± 25.3 | **217.0** ± 21.3 | **275.2** ± 24.7 |
| **General passive behavior** | **257.8** ± 54.6 | **201.8** ± 62.0 | **212.7** ± 57.7 | **328.0** ± 108.0 | **285.8** ± 108.5 | **243.3** ± 109.1 | **167.7** ± 56.0 | **212.1** ± 94.2 | **393.5** ± 51.1 | **281.9** ± 113.6 | **345.8** ± 73.4 | **147.5** ± 45.1 | **193.4** ± 48.5 | **145.5** ± 45.1 | **117.2** ± 55.5 | **30.6** ± 15.4 |
| **Conflict** | **3.9** ± 1.6 | **0.5** ± 0.5 | **2.1** ± 1.0 | **1.1** ± 0.8 | **21.9** ± 15.8 | **60.9** ± 23.9 | **11.4** ± 7.5 | **31.3** ± 19.9 | **4.6** ± 1.8 | **23.0** ± 14.5 | **15.2** ± 7.0 | **13.9** ± 6.0 | **5.8** ± 3.1 | **12.7** ± 6.7 | **43.7** ± 27.4 | **29.5** ± 12.3 |
| **Self-grooming** | **106.4** ± 20.4 | **117.9** ± 37.8 | **130.0** ± 28.3 | **102.2** ± 18.3 | **100.9** ± 35.0 | **92.1** ± 29.8 | **78.7** ± 16.8 | **92.0** ± 38.2 | **113.3** ± 13.3 | **115.5** ± 37.7 | **128.2** ± 23.8 | **144.9** ± 42.1 | **106.3** ± 24.1 | **96.8** ± 22.3 | **176.6** ± 42.0 | **75.2** ± 20.5 |
| **Allogrooming** | **2.2** ± 0.9 | **24.1** ± 7.7 | **3.7** ± 2.1 | **1.6** ± 1.1 | **1.2** ± 1.2 | **1.8** ± 1.8 | **4.2** ± 4.2 | **2.2** ± 2.2 | **6.6** ± 1.9 | **9.1** ± 5.5 | **8.5** ± 3.3 | **0.0** ± 0.0 | **3.1** ± 2.8 | **1.4** ± 1.4 | **7.2** ± 7.2 | **0.7** ± 0.4 |
| **Sniffing others** | **116.8** ± 17.1 | **170.0** ± 32.7 | **122.0** ± 19.9 | **103.0** ± 20.3 | **173.2** ± 42.3 | **204.4** ± 44.9 | **215.1** ± 54.4 | **157.0** ± 61.4 | **86.6** ± 16.7 | **123.1** ± 30.1 | **104.0** ± 12.3 | **125.1** ± 18.7 | **216.8** ± 49.5 | **195.5** ± 25.5 | **162.9** ± 22.6 | **253.2** ± 42.2 |
| **Sniffing anogenitally** | **3.6** ± 2.1 | **9.3** ± 8.2 | **8.6** ± 4.4 | **2.5** ± 2.3 | **7.1** ± 2.4 | **3.5** ± 1.2 | **9.8** ± 4.1 | **7.1** ± 3.6 | **1.8** ± 0.8 | **7.8** ± 6.8 | **1.3** ± 0.7 | **1.3** ± 0.7 | **21.5** ± 11.5 | **12.7** ± 5.9 | **18.6** ± 10.7 | **16.5** ± 7.9 |
| **In opening facing open field** | **49.1** ± 16.9 | **14.8** ± 6.0 | **50.2** ± 10.1 | **36.0** ± 12.0 | **67.9** ± 32.5 | **59.9** ± 24.7 | **79.8** ± 19.2 | **130.2** ± 22.5 | **26.7** ± 4.5 | **13.5** ± 7.2 | **26.2** ± 11.4 | **53.2** ± 21.9 | **76.1** ± 34.1 | **122.8** ± 48.5 | **142.8** ± 49.1 | **193.9** ± 36.4 |
| **Resting/immobile** | **242.8** ± 53.0 | **126.3** ± 35.3 | **192.6** ± 55.5 | **259.1** ± 91.6 | **280.2** ± 109.4 | **237.1** ± 108.8 | **160.9** ± 54.1 | **208.9** ± 94.3 | **310.9** ± 53.5 | **277.0** ± 110.9 | **222.6** ± 47.0 | **129.1** ± 38.6 | **190.0** ± 47.2 | **144.0** ± 45.0 | **110.3** ± 56.0 | **29.2** ± 14.6 |
| **Self-grooming alone** | **103.6** ± 20.2 | **98.7** ± 33.4 | **113.8** ± 24.4 | **91.5** ± 16.1 | **95.8** ± 36.0 | **86.6** ± 27.4 | **78.1** ± 17.1 | **90.5** ± 36.9 | **105.3** ± 14.0 | **109.4** ± 36.9 | **103.9** ± 23.0 | **143.2** ± 41.2 | **94.5** ± 22.9 | **96.2** ± 22.5 | **167.5** ± 37.0 | **74.7** ± 20.5 |
| **Boxing** | **2.7** ± 1.2 | **0.0** ± 0.0 | **1.0** ± 0.5 | **0.0** ± 0.0 | **1.8** ± 1.3 | **1.4** ± 1.0 | **1.0** ± 1.0 | **8.2** ± 8.0 | **2.3** ± 1.5 | **0.4** ± 0.4 | **3.4** ± 1.8 | **4.0** ± 3.6 | **3.2** ± 1.8 | **1.2** ± 1.2 | **5.1** ± 2.4 | **12.9** ± 8.0 |
| **Nose-Off** | **0.4** ± 0.4 | **0.0** ± 0.0 | **0.7** ± 0.4 | **0.8** ± 0.5 | **19.0** ± 15.9 | **41.5** ± 23.6 | **8.3** ± 7.7 | **23.0** ± 20.2 | **0.2** ± 0.1 | **0.0** ± 0.0 | **0.2** ± 0.2 | **0.0** ± 0.0 | **1.8** ± 1.8 | **8.1** ± 5.0 | **21.1** ± 15.6 | **6.3** ± 2.2 |
| **Fighting with other** | **0.0** ± 0.0 | **0.0** ± 0.0 | **0.0** ± 0.0 | **0.0** ± 0.0 | **1.2** ± 0.6 | **1.9** ± 1.1 | **1.2** ± 1.2 | **0.0** ± 0.0 | **0.0** ± 0.0 | **0.4** ± 0.4 | **0.0** ± 0.0 | **1.4** ± 1.4 | **0.3** ± 0.3 | **0.0** ± 0.0 | **0.0** ± 0.0 | **0.0** ± 0.0 |
| **Flee** | **0.0** ± 0.0 | **0.0** ± 0.0 | **0.0** ± 0.0 | **0.0** ± 0.0 | **0.0** ± 0.0 | **0.7** ± 0.6 | **0.5** ± 0.5 | **0.0** ± 0.0 | **0.0** ± 0.0 | **0.0** ± 0.0 | **0.0** ± 0.0 | **0.0** ± 0.0 | **0.2** ± 0.2 | **0.5** ± 0.3 | **0.5** ± 0.5 | **4.4** ± 3.6 |
| **Defensive stance** | **0.0** ± 0.0 | **0.0** ± 0.0 | **0.0** ± 0.0 | **0.0** ± 0.0 | **0.0** ± 0.0 | **14.7** ± 13.3 | **0.5** ± 0.5 | **0.0** ± 0.0 | **0.0** ± 0.0 | **0.0** ± 0.0 | **0.0** ± 0.0 | **0.0** ± 0.0 | **0.3** ± 0.3 | **0.8** ± 0.6 | **13.4** ± 13.4 | **3.3** ± 3.3 |
| **Chasing (after fight)** | **0.0** ± 0.0 | **0.0** ± 0.0 | **0.0** ± 0.0 | **0.0** ± 0.0 | **0.0** ± 0.0 | **0.0** ± 0.0 | **0.0** ± 0.0 | **0.0** ± 0.0 | **0.0** ± 0.0 | **0.0** ± 0.0 | **0.0** ± 0.0 | **0.0** ± 0.0 | **0.0** ± 0.0 | **0.0** ± 0.0 | **0.0** ± 0.0 | **0.0** ± 0.0 |
| **Wrestling** | **0.8** ± 0.7 | **0.5** ± 0.5 | **0.5** ± 0.5 | **0.3** ± 0.3 | **0.0** ± 0.0 | **0.7** ± 0.7 | **0.0** ± 0.0 | **0.0** ± 0.0 | **2.1** ± 0.9 | **22.2** ± 14.1 | **11.5** ± 5.6 | **8.5** ± 4.0 | **0.0** ± 0.0 | **1.9** ± 1.9 | **3.5** ± 2.7 | **2.5** ± 2.5 |
| **Self-grooming near others** | **2.8** ± 2.3 | **19.2** ± 10.4 | **16.2** ± 5.4 | **10.7** ± 6.3 | **5.1** ± 3.4 | **5.5** ± 5.5 | **0.6** ± 0.6 | **1.5** ± 1.5 | **8.0** ± 4.6 | **6.1** ± 3.0 | **24.3** ± 13.8 | **1.7** ± 1.4 | **11.8** ± 6.7 | **0.7** ± 0.7 | **9.0** ± 9.0 | **0.5** ± 0.5 |
| **In opening facing burrow** | **0.3** ± 0.2 | **0.0** ± 0.0 | **0.4** ± 0.4 | **0.0** ± 0.0 | **0.1** ± 0.1 | **0.1** ± 0.1 | **0.7** ± 0.4 | **2.2** ± 1.4 | **0.0** ± 0.0 | **0.2** ± 0.1 | **0.1** ± 0.1 | **0.2** ± 0.2 | **0.0** ± 0.0 | **0.1** ± 0.1 | **0.0** ± 0.0 | **0.5** ± 0.4 |
| **Eating** | **4.1** ± 1.8 | **0.0** ± 0.0 | **12.7** ± 12.1 | **12.0** ± 9.6 | **0.0** ± 0.0 | **1.4** ± 1.4 | **0.0** ± 0.0 | **8.0** ± 8.0 | **0.1** ± 0.1 | **0.0** ± 0.0 | **4.1** ± 2.5 | **0.1** ± 0.1 | **1.3** ± 1.3 | **0.0** ± 0.0 | **0.0** ± 0.0 | **0.0** ± 0.0 |
| **Digging/moving bedding** | **43.3** ± 9.9 | **29.7** ± 12.4 | **33.4** ± 9.5 | **42.3** ± 17.7 | **6.9** ± 4.5 | **6.2** ± 4.6 | **39.0** ± 33.7 | **4.2** ± 3.9 | **47.3** ± 12.5 | **44.0** ± 15.3 | **44.8** ± 14.3 | **42.5** ± 13.8 | **6.5** ± 4.3 | **4.1** ± 3.8 | **3.2** ± 2.2 | **9.0** ± 3.9 |
| **Drinking** | **8.0** ± 3.8 | **4.9** ± 2.7 | **19.0** ± 4.5 | **15.2** ± 5.7 | **0.7** ± 0.7 | **3.6** ± 3.2 | **11.0** ± 7.6 | **2.8** ± 2.8 | **14.6** ± 6.8 | **8.8** ± 5.1 | **13.2** ± 4.8 | **13.7** ± 6.4 | **28.4** ± 16.4 | **23.4** ± 18.3 | **10.9** ± 5.6 | **15.9** ± 9.6 |
| **Walking** | **261.4** ± 37.3 | **288.0** ± 48.9 | **261.2** ± 33.8 | **225.9** ± 46.0 | **225.9** ± 55.9 | **211.5** ± 42.0 | **262.0** ± 28.2 | **234.4** ± 44.5 | **173.4** ± 22.0 | **241.4** ± 62.4 | **183.8** ± 28.4 | **314.4** ± 37.7 | **234.1** ± 30.5 | **275.4** ± 24.0 | **208.3** ± 20.9 | **271.3** ± 24.8 |
| **Walk over/under others** | **1.8** ± 0.9 | **7.3** ± 3.8 | **3.0** ± 0.8 | **2.8** ± 1.3 | **1.4** ± 1.2 | **2.2** ± 1.6 | **1.0** ± 0.4 | **0.0** ± 0.0 | **1.5** ± 0.6 | **0.5** ± 0.3 | **1.4** ± 0.7 | **4.3** ± 4.3 | **1.7** ± 1.1 | **1.6** ± 0.5 | **2.1** ± 1.8 | **0.3** ± 0.3 |
| **Non-social exploration** | **41.7** ± 5.1 | **31.1** ± 9.4 | **41.5** ± 8.2 | **28.1** ± 12.0 | **7.0** ± 3.7 | **9.2** ± 5.8 | **20.2** ± 8.3 | **16.8** ± 15.6 | **27.2** ± 6.0 | **31.4** ± 8.0 | **23.2** ± 7.2 | **39.2** ± 13.5 | **5.0** ± 2.3 | **8.0** ± 5.0 | **6.5** ± 5.6 | **3.6** ± 1.3 |
| Episodes of | | | | | | | | | | | | | | | | |
| **Active social behavior** | **18.6** ± 2.4 | **23.4** ± 3.6 | **19.9** ± 2.8 | **15.3** ± 3.6 | **23.1** ± 6.4 | **25.2** ± 6.0 | **26.2** ± 3.5 | **29.2** ± 12.1 | **14.2** ± 2.4 | **31.5** ± 11.5 | **18.5** ± 2.9 | **28.0** ± 5.3 | **38.3** ± 8.4 | **46.4** ± 11.0 | **34.8** ± 6.2 | **54.7** ± 10.4 |
| **Passive Social behavior** | **1.0** ± 0.3 | **2.0** ± 0.7 | **0.7** ± 0.2 | **0.8** ± 0.3 | **0.8** ± 0.4 | **0.6** ± 0.3 | **0.5** ± 0.2 | **0.4** ± 0.4 | **0.8** ± 0.2 | **0.6** ± 0.4 | **0.7** ± 0.2 | **1.1** ± 0.6 | **0.4** ± 0.3 | **0.1** ± 0.1 | **0.1** ± 0.1 | **0.1** ± 0.1 |
| **Social context** | **19.8** ± 2.5 | **26.1** ± 3.5 | **21.4** ± 2.7 | **16.6** ± 3.3 | **24.3** ± 6.7 | **25.9** ± 5.9 | **26.8** ± 3.3 | **29.8** ± 11.9 | **15.4** ± 2.4 | **32.4** ± 11.4 | **19.8** ± 2.8 | **29.4** ± 5.4 | **39.6** ± 8.2 | **46.6** ± 11.0 | **35.2** ± 6.4 | **54.9** ± 10.4 |
| **General active behavior** | **29.3** ± 3.0 | **29.1** ± 5.6 | **30.4** ± 3.5 | **24.2** ± 6.2 | **23.2** ± 5.7 | **23.7** ± 5.9 | **30.8** ± 3.6 | **32.8** ± 11.2 | **19.3** ± 2.7 | **35.4** ± 14.0 | **22.1** ± 2.8 | **37.2** ± 4.5 | **31.8** ± 5.9 | **47.3** ± 14.7 | **34.2** ± 3.2 | **53.0** ± 7.3 |
| **General passive behavior** | **6.2** ± 0.7 | **6.1** ± 1.0 | **5.9** ± 0.9 | **3.9** ± 0.6 | **6.9** ± 1.1 | **3.9** ± 1.0 | **4.7** ± 1.1 | **5.0** ± 1.9 | **4.7** ± 0.6 | **3.8** ± 1.3 | **4.3** ± 0.6 | **4.5** ± 1.0 | **5.8** ± 2.2 | **6.0** ± 1.5 | **3.8** ± 1.2 | **1.0** ± 0.5 |
| **Conflict** | **0.6** ± 0.3 | **0.0** ± 0.0 | **0.5** ± 0.3 | **0.2** ± 0.2 | **1.9** ± 0.9 | **3.0** ± 0.8 | **1.0** ± 0.4 | **1.2** ± 0.5 | **0.6** ± 0.2 | **1.6** ± 0.9 | **1.1** ± 0.4 | **1.4** ± 0.5 | **1.3** ± 0.6 | **1.8** ± 0.5 | **2.3** ± 0.4 | **4.2** ± 1.8 |
| **Self-grooming** | **4.4** ± 0.6 | **4.5** ± 1.2 | **4.7** ± 0.5 | **4.0** ± 0.6 | **3.4** ± 0.7 | **3.0** ± 0.9 | **4.2** ± 0.5 | **4.0** ± 1.5 | **4.6** ± 0.6 | **2.6** ± 0.7 | **4.4** ± 0.5 | **5.3** ± 1.4 | **5.6** ± 0.7 | **4.7** ± 1.0 | **6.9** ± 1.5 | **3.5** ± 0.5 |
| **Allogrooming** | **0.3** ± 0.2 | **1.1** ± 0.4 | **0.3** ± 0.2 | **0.1** ± 0.1 | **0.1** ± 0.1 | **0.1** ± 0.1 | **0.0** ± 0.0 | **0.2** ± 0.2 | **0.4** ± 0.1 | **0.7** ± 0.4 | **0.4** ± 0.1 | **0.0** ± 0.0 | **0.1** ± 0.1 | **0.2** ± 0.2 | **0.7** ± 0.7 | **0.1** ± 0.1 |
| **Sniffing others** | **17.7** ± 2.1 | **21.1** ± 3.6 | **18.0** ± 2.2 | **14.8** ± 3.5 | **21.8** ± 6.3 | **24.4** ± 6.0 | **25.3** ± 3.3 | **27.5** ± 11.5 | **13.7** ± 2.3 | **29.8** ± 11.1 | **17.9** ± 3.0 | **27.5** ± 5.0 | **35.5** ± 7.7 | **44.1** ± 10.4 | **31.7** ± 5.2 | **52.3** ± 9.5 |
| **Sniffing anogenitally** | **0.6** ± 0.3 | **1.1** ± 0.9 | **1.6** ± 0.9 | **0.3** ± 0.3 | **1.2** ± 0.4 | **0.7** ± 0.2 | **0.8** ± 0.3 | **1.5** ± 0.7 | **0.2** ± 0.1 | **1.0** ± 0.6 | **0.2** ± 0.1 | **0.6** ± 0.3 | **2.7** ± 1.4 | **2.1** ± 0.8 | **2.4** ± 1.2 | **2.2** ± 1.1 |
| **Fighting with other** | **0.0** ± 0.0 | **0.0** ± 0.0 | **0.0** ± 0.0 | **0.0** ± 0.0 | **0.4** ± 0.2 | **0.4** ± 0.2 | **0.2** ± 0.2 | **0.0** ± 0.0 | **0.0** ± 0.0 | **0.1** ± 0.1 | **0.0** ± 0.0 | **0.1** ± 0.1 | **0.2** ± 0.2 | **0.0** ± 0.0 | **0.0** ± 0.0 | **0.0** ± 0.0 |
| **Boxing** | **0.5** ± 0.2 | **0.0** ± 0.0 | **0.3** ± 0.2 | **0.0** ± 0.0 | **0.4** ± 0.2 | **0.3** ± 0.2 | **0.1** ± 0.1 | **0.6** ± 0.4 | **0.3** ± 0.2 | **0.1** ± 0.1 | **0.4** ± 0.1 | **0.5** ± 0.4 | **0.8** ± 0.3 | **0.3** ± 0.2 | **0.9** ± 0.3 | **1.1** ± 0.6 |
| **Nose-Off** | **0.1** ± 0.1 | **0.0** ± 0.0 | **0.2** ± 0.1 | **0.2** ± 0.1 | **0.9** ± 0.7 | **1.5** ± 0.5 | **0.3** ± 0.2 | **0.6** ± 0.3 | **0.1** ± 0.0 | **0.0** ± 0.0 | **0.1** ± 0.1 | **0.0** ± 0.0 | **0.2** ± 0.2 | **0.8** ± 0.2 | **0.6** ± 0.3 | **1.2** ± 0.5 |

Supplementary Table S1

The data represents average time or average number of episodes performing named behaviour +/-SEM.

Differences compared to CTR-rats are highlighted where Mann Whitney U p<0.05

Differences compared to non-witnesses are highlighted where Mann Whitney U p<0.05

| | Baseline | | | | | | | |
|---|---|---|---|---|---|---|---|---|
| | Female | | | | Male | | | |
| | CTR | | FLX | | CTR | | FLX | |
| | Non-Witness | Witness | Non-Witness | Witness | Non-Witness | Witness | Non-Witness | Witness |
| n | 8 | 7 | 8 | 6 | 7 | 7 | 6 | 5 |
| Flee | 0.0 ± 0.0 | 0.0 ± 0.0 | 0.0 ± 0.0 | 0.0 ± 0.0 | 0.0 ± 0.0 | 0.4 ± 0.3 | 0.1 ± 0.1 | 0.0 ± 0.0 |
| Wrestling | 0.1 ± 0.0 | 0.0 ± 0.0 | 0.0 ± 0.0 | 0.1 ± 0.1 | 0.0 ± 0.0 | 0.1 ± 0.1 | 0.0 ± 0.0 | 0.0 ± 0.0 |
| Chasing (after fight) | 0.0 ± 0.0 | 0.0 ± 0.0 | 0.0 ± 0.0 | 0.0 ± 0.0 | 0.0 ± 0.0 | 0.0 ± 0.0 | 0.0 ± 0.0 | 0.0 ± 0.0 |
| Defensive stance | 0.0 ± 0.0 | 0.0 ± 0.0 | 0.0 ± 0.0 | 0.0 ± 0.0 | 0.0 ± 0.0 | 0.4 ± 0.2 | 0.1 ± 0.1 | 0.0 ± 0.0 |
| Self-grooming alone | 4.3 ± 0.6 | 3.8 ± 1.0 | 4.0 ± 0.4 | 3.5 ± 0.6 | 3.0 ± 0.7 | 2.9 ± 0.8 | 4.1 ± 0.5 | 3.8 ± 1.4 |
| In opening facing open field | 5.1 ± 0.9 | 2.1 ± 0.6 | 5.3 ± 0.8 | 4.3 ± 1.2 | 9.6 ± 4.6 | 8.9 ± 4.0 | 8.4 ± 2.0 | 11.8 ± 3.8 |
| Eating | 0.4 ± 0.2 | 0.0 ± 0.0 | 0.1 ± 0.0 | 0.3 ± 0.3 | 0.0 ± 0.0 | 0.1 ± 0.1 | 0.0 ± 0.0 | 0.1 ± 0.1 |
| Digging/moving bedding | 2.7 ± 0.8 | 2.0 ± 0.7 | 1.8 ± 0.6 | 2.2 ± 0.8 | 0.6 ± 0.6 | 0.3 ± 0.2 | 2.9 ± 1.8 | 0.6 ± 0.5 |
| Drinking | 1.0 ± 0.4 | 0.6 ± 0.3 | 1.6 ± 0.2 | 1.6 ± 0.6 | 0.1 ± 0.1 | 0.2 ± 0.1 | 0.5 ± 0.3 | 0.3 ± 0.3 |
| Walk over/under others | 0.6 ± 0.3 | 2.0 ± 0.9 | 0.9 ± 0.3 | 0.5 ± 0.2 | 0.1 ± 0.1 | 0.5 ± 0.3 | 0.4 ± 0.2 | 0.0 ± 0.0 |
| Walking | 24.9 ± 2.8 | 24.3 ± 5.2 | 25.9 ± 3.4 | 21.2 ± 5.6 | 21.9 ± 5.7 | 22.2 ± 6.1 | 28.8 ± 3.6 | 32.0 ± 11.4 |
| Resting/immobile | 5.1 ± 0.6 | 4.1 ± 0.6 | 5.2 ± 0.8 | 3.2 ± 0.5 | 6.1 ± 0.9 | 3.4 ± 1.0 | 4.2 ± 1.0 | 4.6 ± 1.7 |
| In opening facing burrow | 0.1 ± 0.1 | 0.0 ± 0.0 | 0.1 ± 0.1 | 0.0 ± 0.0 | 0.1 ± 0.1 | 0.0 ± 0.0 | 0.3 ± 0.2 | 0.3 ± 0.2 |
| Self-grooming near others | 0.1 ± 0.1 | 0.7 ± 0.3 | 0.7 ± 0.2 | 0.6 ± 0.2 | 0.4 ± 0.3 | 0.1 ± 0.1 | 0.1 ± 0.1 | 0.2 ± 0.2 |
| Non-social exploration | 3.8 ± 0.6 | 2.7 ± 0.6 | 3.5 ± 0.6 | 2.5 ± 0.9 | 1.2 ± 0.6 | 1.0 ± 0.5 | 1.6 ± 0.6 | 0.8 ± 0.6 |

| | Post-stress | | | | | | | |
|---|---|---|---|---|---|---|---|---|
| | Female | | | | Male | | | |
| | CTR | | FLX | | CTR | | FLX | |
| | Non-Witness | Witness | Non-Witness | Witness | Non-Witness | Witness | Non-Witness | Witness |
| n | 8 | 7 | 8 | 7 | 6 | 6 | 6 | 5 |
| Flee | 0.0 ± 0.0 | 0.0 ± 0.0 | 0.0 ± 0.0 | 0.0 ± 0.0 | 0.2 ± 0.1 | 0.3 ± 0.2 | 0.3 ± 0.3 | 1.3 ± 0.7 |
| Wrestling | 0.2 ± 0.1 | 1.5 ± 0.9 | 0.6 ± 0.3 | 0.8 ± 0.2 | 0.0 ± 0.0 | 0.1 ± 0.1 | 0.3 ± 0.2 | 0.2 ± 0.2 |
| Chasing (after fight) | 0.0 ± 0.0 | 0.0 ± 0.0 | 0.0 ± 0.0 | 0.0 ± 0.0 | 0.0 ± 0.0 | 0.0 ± 0.0 | 0.0 ± 0.0 | 0.0 ± 0.0 |
| Defensive stance | 0.0 ± 0.0 | 0.0 ± 0.0 | 0.0 ± 0.0 | 0.0 ± 0.0 | 0.1 ± 0.1 | 0.3 ± 0.2 | 0.2 ± 0.2 | 0.4 ± 0.4 |
| Self-grooming alone | 4.2 ± 0.6 | 2.3 ± 0.6 | 3.8 ± 0.4 | 5.0 ± 1.4 | 4.7 ± 0.5 | 4.6 ± 1.0 | 6.5 ± 1.3 | 3.4 ± 0.5 |
| In opening facing open field | 3.0 ± 0.5 | 2.4 ± 1.4 | 3.3 ± 1.1 | 7.5 ± 2.4 | 11.1 ± 5.0 | 19.4 ± 9.4 | 12.7 ± 2.5 | 25.8 ± 4.3 |
| Eating | 0.0 ± 0.0 | 0.0 ± 0.0 | 0.2 ± 0.1 | 0.0 ± 0.0 | 0.2 ± 0.2 | 0.0 ± 0.0 | 0.0 ± 0.0 | 0.0 ± 0.0 |
| Digging/moving bedding | 3.0 ± 1.0 | 4.1 ± 1.8 | 3.1 ± 0.8 | 4.0 ± 1.1 | 0.6 ± 0.2 | 0.6 ± 0.6 | 0.3 ± 0.2 | 1.2 ± 0.4 |
| Drinking | 0.9 ± 0.3 | 1.1 ± 0.6 | 1.1 ± 0.3 | 1.0 ± 0.7 | 0.9 ± 0.3 | 0.6 ± 0.3 | 1.0 ± 0.6 | 1.7 ± 0.9 |
| Walk over/under others | 0.4 ± 0.2 | 0.4 ± 0.2 | 0.4 ± 0.2 | 1.0 ± 1.0 | 0.5 ± 0.3 | 0.8 ± 0.3 | 0.5 ± 0.3 | 0.1 ± 0.1 |
| Walking | 16.6 ± 2.7 | 32.8 ± 13.4 | 19.6 ± 2.6 | 34.0 ± 3.8 | 30.4 ± 5.8 | 45.8 ± 14.5 | 33.2 ± 3.5 | 52.1 ± 7.4 |
| Resting/immobile | 3.9 ± 0.7 | 3.2 ± 1.2 | 3.6 ± 0.5 | 3.5 ± 0.8 | 5.3 ± 2.1 | 5.9 ± 1.5 | 3.7 ± 1.2 | 0.9 ± 0.4 |
| In opening facing burrow | 0.0 ± 0.0 | 0.2 ± 0.1 | 0.0 ± 0.0 | 0.1 ± 0.1 | 0.0 ± 0.0 | 0.2 ± 0.2 | 0.0 ± 0.0 | 0.2 ± 0.1 |
| Self-grooming near others | 0.4 ± 0.1 | 0.4 ± 0.2 | 0.6 ± 0.2 | 0.3 ± 0.2 | 0.9 ± 0.5 | 0.1 ± 0.1 | 0.3 ± 0.3 | 0.1 ± 0.1 |
| Non-social exploration | 2.3 ± 0.4 | 2.3 ± 0.7 | 2.2 ± 0.5 | 2.2 ± 0.6 | 0.9 ± 0.3 | 0.8 ± 0.3 | 0.5 ± 0.4 | 0.8 ± 0.2 |
